# Treacle’s ability to form liquid-like phase condensates is essential for nucleolar fibrillar center assembly, efficient rRNA transcription and processing, and rRNA gene repair

**DOI:** 10.1101/2023.05.11.540311

**Authors:** Artem K. Velichko, Nadezhda V. Petrova, Dmitry A. Deriglazov, Anastasia P. Kovina, Artem V. Luzhin, Eugene P. Kazakov, Igor I. Kireev, Sergey V. Razin, Omar L. Kantidze

## Abstract

We investigated the role of the nucleolar protein Treacle in organizing and regulating the nucleolus in human cells. Our results support Treacle’s ability to form liquid-like phase condensates through electrostatic interactions among molecules. The formation of these biomolecular condensates is crucial for segregating nucleolar fibrillar centers from the dense fibrillar component and ensuring high levels of rRNA gene transcription and accurate rRNA processing. Both the central and C-terminal domains of Treacle are required to form liquid-like condensates. The initiation of phase separation is attributed to the C-terminal domain. The central domain is characterized by repeated stretches of alternatively charged amino-acid residues and is vital for condensate stability. Overexpression of mutant forms of Treacle that cannot form liquid-like phase condensates compromises the assembly of fibrillar centers, suppressing rRNA gene transcription and disrupting rRNA processing. These mutant forms also fail to recruit DNA topoisomerase II binding protein 1 (TOPBP1), suppressing the DNA damage response in the nucleolus.

## Introduction

Applying polymer chemistry principles to biological molecules has significantly broadened our understanding of the functions of membraneless compartments (Mehta & Zhang, 2022; Mittag & Pappu, 2022). Multiple weak, cooperative, and dynamic interactions underlie the assembly of these self-organized structures, termed biomolecular condensates. Mechanistically, liquid–liquid phase separation (LLPS)—a demixing process that yields a condensed protein-enriched phase and a dilute phase—can mediate the self-assembly of biomolecular condensates. In biological systems, phase separation is facilitated by a combination of multivalent interactions mediated by intrinsically disordered regions and site-specific interactions that drive percolation (Banani et al., 2017; Shin & Brangwynne, 2017; Uversky, 2019). Biomolecular condensates serve as functional hubs in diverse cellular processes such as transcription (Sabari et al., 2020), microtubule nucleation (Woodruff et al., 2017), and the adaptive stress response (Franzmann & Alberti, 2019).

The nucleolus is a large and complex subnuclear compartment that can also be considered a multicomponent and multilayered biomolecular condensate (Lafontaine et al., 2021). It is formed around arrays of ribosomal gene repeats (rDNA), which are transcribed by RNA polymerase I (RNA Pol I) to produce ribosomal RNA (rRNA) (Cerqueira & Lemos, 2019). In the nucleolus, RNAs and hundreds of different proteins are segregated into three immiscible phases: the fibrillar center (FC), which is the RNA Pol I transcription factory; the dense fibrillar component (DFC), which is the intermediate layer rich in fibrillarin (FBL); and the granular component (GC), which is the outer layer rich in nucleophosmin (NPM1/B23). The mechanisms underlying DFC and GC formation through phase separation are relatively well understood. Studies have demonstrated that a ternary mixture comprising the GC protein NPM1, a monomeric version of the DFC protein FBL, and generic rRNA is sufficient to form an FBL-enriched inner phase that coexists with an NPM-enriched outer phase (Feric et al., 2016; Mitrea et al., 2016; Mitrea et al., 2018). In contrast, the mechanisms of FC formation and the role of LLPS in this process have received considerably less attention.

Previous studies on FC organization in living cells suggest that FC assembly may also occur via phase separation (Falahati et al., 2016; Falahati & Wieschaus, 2017). Several lines of evidence support this hypothesis. Firstly, FC components migrate to the nucleolar periphery and aggregate into large structures known as nucleolar caps when RNA Pol I transcription is inhibited by actinomycin D (AMD) or genotoxic drugs (Harding et al., 2015; Korsholm et al., 2019; Reynolds et al., 1964). This structural transformation is reminiscent of liquid droplet fusion. Secondly, high-speed molecular dynamics indicative of fluid-like behavior have been observed in nucleolar caps using single-molecule tracking of RNA Pol I and chromatin-bound upstream binding transcription factor (UBFT/UBF) (Ide et al., 2020). Thirdly, both UBF and its partner treacle ribosome biogenesis factor 1 (TCOF1/Treacle) demonstrated the ability to undergo condensation *in vitro* and *in vivo*, respectively (Jaberi-Lashkari et al., 2023; King et al., 2024). Therefore, it is likely that FC assembly and organization rely on LLPS, similar to other nucleolar subcompartments. However, further studies are needed to elucidate the molecular determinants and biophysical properties underlying this process.

In this context, the nucleolar phosphoprotein Treacle is of particular interest. As previously mentioned, it directly interacts with UBF and RNA Pol I, colocalizing with them within the FC. In addition, Treacle exhibits both transcription-dependent and transcription-independent functions (Gal et al., 2022). Its role in ribosome biogenesis is well-documented since it assists in the transcription and processing of rRNA (Gonzales et al., 2005; Lin & Yeh, 2009; Valdez et al., 2004). It is also a master regulator of the DNA damage response (DDR) in the nucleolus, mediating the recruitment of repair factors to rDNA damaged by I-PpoI-induced breaks, replication stress, or R-loop formation (Ciccia et al., 2014; Korsholm et al., 2019; Larsen et al., 2014; Mooser et al., 2020; Velichko et al., 2019, 2021). Therefore, it acts as a nucleolar hub with both transcriptional and DNA repair functions.

The multifunctionality of Treacle may be related to its ability to engage in multivalent interactions and undergo phase separation because of the presence of extended intrinsically disordered regions. In addition, Treacle’s partners in transcription (RNA Pol I, UBF) and DNA repair (DNA topoisomerase II binding protein 1 [TOPBP1]) also utilize phase separation as a functioning mechanism (Frattini et al., 2021; Ide et al., 2020; King et al., 2024). Transcription of rDNA can be regulated by phase separation of RNA Pol I, while TOPBP1 condensation can activate the ATR serine/threonine kinase (ATR) signaling pathway (Frattini et al., 2021; Ide et al., 2020).

Considering these observations, we conducted an in-depth study of Treacle’s condensation characteristics and their impact on the structural organization and functional dynamics of nucleoli. We found that Treacle can form biomolecular condensates and characterized its structural features regulating this process. Through its ability to condense, Treacle promotes the formation of nucleolar FCs by recruiting and concentrating transcription factors at rDNA and separating FC components from the DFC. Collectively, this segregates rRNA synthesis and subsequent processing in distinct compartments of the nucleolus. Impairing Treacle’s condensation ability results in the mixing of FC and DFC components, reducing the efficiency of both processes, equivalent to completed depleting endogenous Treacle. We also demonstrate that Treacle’s phase separation is critical for its interaction with TOPBP1 and activation of the DDR in rDNA during genotoxic stress. Our findings reveal the role of Treacle not only as a structural scaffold for FCs but also as a nucleolar hub integrating the functions of rDNA transcription, rRNA processing, and preserving rDNA integrity.

### Treacle is a scaffold protein for nucleolar FCs and DFCs

First, we defined the role of Treacle in the structural organization of the nucleolus. We employed the CRISPR/Cas9 system to knock out the *TCOF1* gene encoding Treacle in HeLa cells to obtain a population of cells lacking Treacle signals (Treacle-negative cells; Fig. S1A). Immunocytochemical staining for UBF and RNA Pol I subunit A (POLR1A/RPA194; FC markers), FBL (DFC marker), and B23 and nucleolin (NCL; GC markers) revealed that Treacle-positive cells exhibited a classical tripartite nucleolar structure, with Treacle colocalizing with UBF1 and RPA194 in FCs, FBL in DFCs, and B23 and NCL surrounding DFCs (Fig. 1A). Treacle depletion resulted in the disappearance of distinct FC and DFC structures, causing RPA194, UBF, and FBL to diffuse throughout the nucleolus (Figs. 1B, S1A). This suggests a mixing of FC and DFC components (Fig. S1B). The organization of the GC component, however, was minimally affected (Figs. 1B, S1C). Notably, Treacle depletion also partially redistributed RPA194, UBF, FBL, and NCL from the nucleolus to the nucleoplasm, likely reflecting disrupted ribosomal gene transcription (Figs. 1B, S1A). Similar effects were observed in normal human skin fibroblasts following Treacle depletion (Fig. S1D–E). These results indicated that Treacle plays a critical role in forming the integrated structure of FCs and DFCs.

**Fig. 1.**
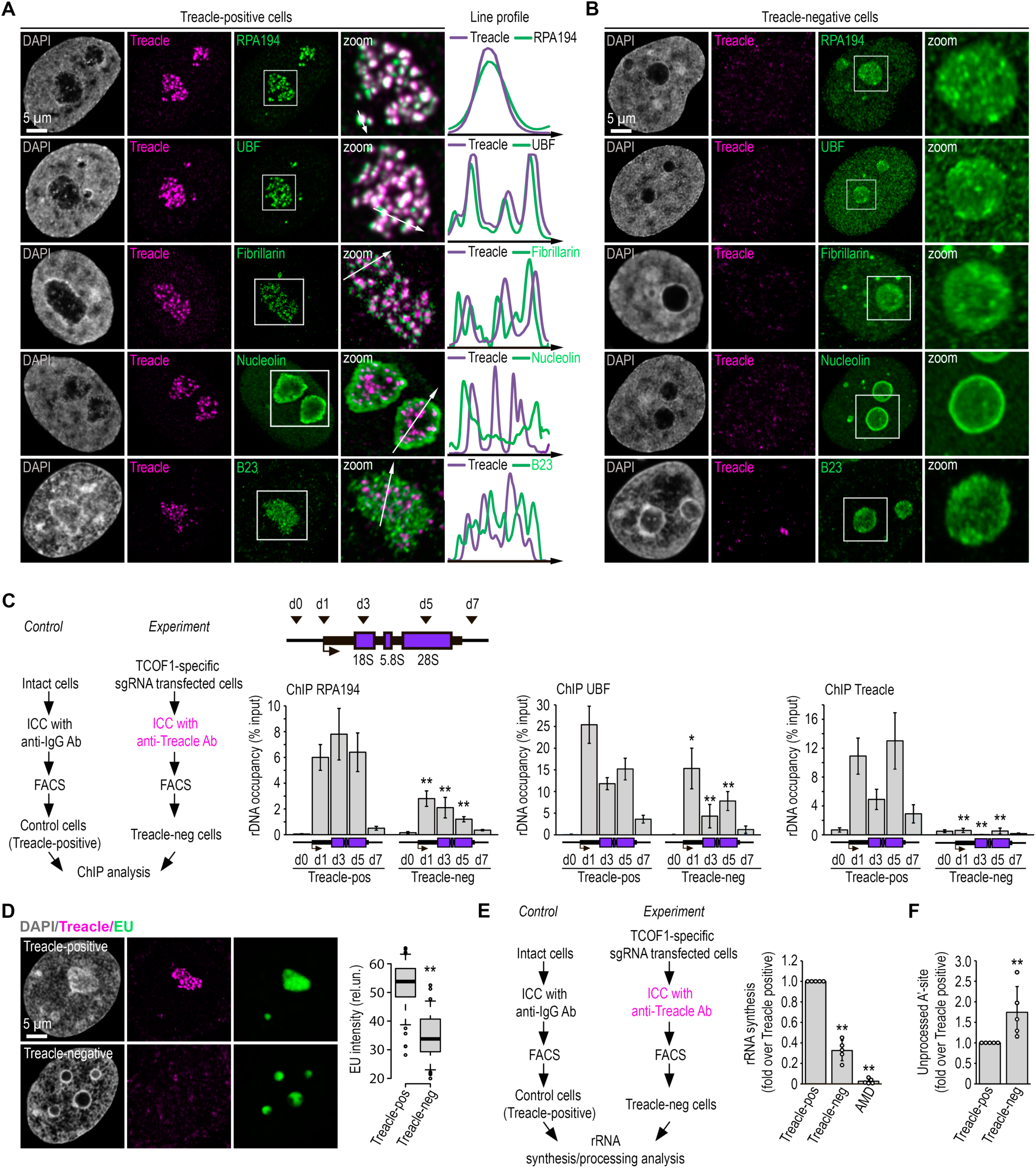
Treacle is a scaffold protein for nucleolar FCs and DFCs. **(A)** Intact HeLa cells (Treacle-positive) were fixed and co-immunostained with Treacle and with either RPA194, UBF, Fibrillarin, B23 or Nucleolin antibodies. DNA was stained with DAPI (gray). Cells were analyzed by laser scanning confocal microscopy. Representative images of cells and nucleoli (magnified images) are shown. Co-localization analysis was performed on the merged images. Graphs illustrate quantification in arbitrary units of fluorescence distribution along the lines shown in the figures. **(B)** HeLa cells were transfected with a construct coding CRISPR/Cas9 and sgRNA to the TCOF1 gene. After 7-10 days after transfection, the cells were fixed and co-immunostained with Treacle and either RPA194, UBF, Fibrillarin, B23 or Nucleolin antibodies. DNA was stained with DAPI (gray). Cells were analyzed by laser scanning confocal microscopy. Representative images of Treacle-negative cells and nucleoli (magnified images) are shown. **(C)** HeLa cells were transfected with a construct coding CRISPR/Cas9 and sgRNA to the TCOF1 gene (*experiment*). After 7-10 days after transfection cell were fixed, immunostained with Treacle antibodies and subjected to cell sorting in the fluorescent analysis mode to obtain Treacle-negative populations. Intact HeLa cells (*control*) were fixed, immunostained with IgG antibodies, passed through all FACS-related procedure in the light scattering analysis mode and used as a control (Treacle-positive). The sorted cell fractions were used for chromatin immunoprecipitation (ChIP) analysis with either Treacle, RPA194 or UBF antibodies. ChIP was followed by qPCR using the d0, d1, d3, d5, d7 primers to the rRNA gene (positioned as indicated on the scheme). Data are represented relative to the input. Values are means ±SD from at least three independent replicates (**, p < 0.01; *, p < 0.1; by unpaired t test). **(D)** HeLa cells were transfected with a construct coding CRISPR/Cas9 and sgRNA to the TCOF1 gene. After 7-10 days after transfection the cells were pulsed with EU (100 μM for 2 hr), fixed and immunostained with Treacle antibodies. EU (green) was revealed by click chemistry. The DNA was stained with DAPI (gray). Representative images of Treacle-positive and Treacle-negative HeLa cells are shown. Quantification of EU fluorescence intensities in Treacle-positive and Treacle-negative HeLa cells are show in the right panel (**, p < 0.01 by unpaired t test; n>500). **(E)** HeLa cells were transfected with a construct coding CRISPR/Cas9 and sgRNA to the TCOF1 gene (*experiment*). After 7-10 days after transfection the cells were fixed, immunostained with Treacle antibodies and subjected to cell sorting in the fluorescent analysis mode to obtain Treacle-negative populations. Intact HeLa cells (*control*) were fixed, stained with IgG antibodies, passed through all FACS-related procedure in the light scattering analysis mode and used as a control (Treacle-positive). The sorted cell fractions were used for RNA extraction. RT-qPCR was performed; it shows levels of 47S pre-rRNA normalized to GAPDH mRNA. Normalized pre-rRNA level in Treacle-positive cells is set to 1. Values are mean ± SD. The calculation is presented for 5 biological replicates (**, p < 0.01 by unpaired t test). **(F)** Hela cells were processed as described in (E). It shows levels of A’-site contained unprocessed rRNA normalized to GAPDH mRNA. Normalized unprocessed rRNA level in Treacle-positive cells is set to 1. Values are mean ± SD. The calculation is presented for 5 biological replicates (**, p < 0.01 by unpaired t test).

Next, we investigated the impact of nucleolar FC disintegration on the efficiency of rDNA transcription and the recruitment of the RNA Pol I transcription machinery to rDNA. Chromatin immunoprecipitation analysis (ChIP) of cells transfected with an anti-Treacle single guide RNA (sgRNA) revealed a decreased occupancy of UBF and RPA194 within the rDNA coding region (Fig. 1C). In contrast, transfection with a mock sgRNA did not cause such an effect (Fig. S2A). These observations suggest that Treacle amplifies the recruitment rather than initiates the primary binding of the transcription machinery to rDNA. Depleting Treacle substantially suppressed rDNA transcription, although not completely, unlike the repression induced by AMD (Figs. 1D–E, S2B). Furthermore, cells depleted of Treacle exhibited impaired 5′ external transcribed spacer processing, particularly in the cleavage of the A′ site (Fig. 1F), which is one of the initial steps in 18S rRNA maturation.

In summary, Treacle functions as a scaffold protein in nucleolar FCs, and its depletion compromises both rRNA gene transcription and processing.

### Treacle drives the formation of biomolecular condensates

Currently, it has become evident that various scaffold proteins can self-organize by forming liquid-phase condensates (Banani et al., 2017; Shin & Brangwynne, 2017). According to protein structure predictors (e.g., AlphaFold, IUPred2, PONDR, and FuzDrop), Treacle is a fully intrinsically disordered protein (Fig. S3A). Nevertheless, Treacle can be broadly categorized into three distinct functional domains: the N-terminal domain (ND), the central domain, and the C-terminal domain (CD) (Gal et al., 2022). The ND of Treacle harbors a LisH motif and a nuclear localization signal (NLS). The central domain contains 15 low-complexity regions (LCRs) enriched in serine and glutamic acid residues (S/E-rich LCRs), which are interspersed with unstructured linker sequences (Jaberi-Lashkari et al., 2023). The 11 linkers within the central domain and their adjacent S/E-rich LCRs exhibit high homology. Consequently, the central domain will be referred to as the repeating domain (RD) henceforth. The CD harbors additional NLSs, a UBF binding site, and a lysine-rich LCR (K-rich LCR) at the terminus, which also functions as a nucleolar localization signal (NoLS) (Fig. 2A).

**Fig. 2.**
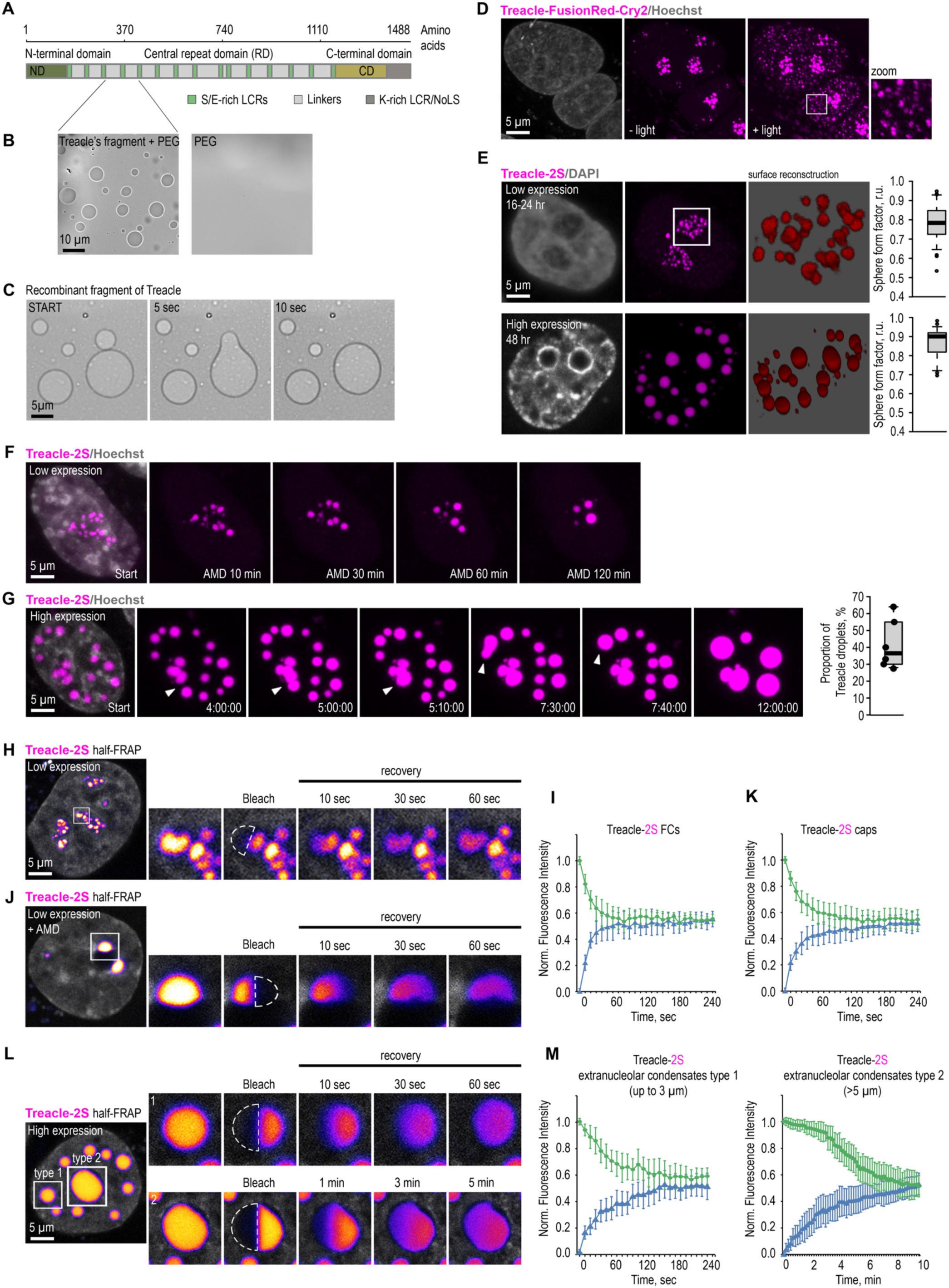
Treacle drives the formation of nuclear condensates. **(A)** The structure of the most common isoform of Treacle. Treacle isoform d (1488 amino acids, 152 kDa, NP_001128715.1) is encoded by the Treacle transcript variant 4. It is an intrinsically disordered protein with N-terminal (ND, 1–83 aa), C-terminal (CD, 1121–1488 aa) regions, and central repeated domain (RD, 83–1121 aa) consisting of 15 low complexity regions (LCR) interspersed with disordered linker sequences. **(B)** The purified recombinant fragment of Treacle undergoes condensation *in vitro* in the presence of 5% polyethylene glycol. **(C)** Condensates of recombinant Treacle’s fragment were time-lapse imaged. Representative images of condensates fusion events are shown. **(D)** HeLa cells were transfected with Treacle-FusionRed-Cry2 (opto-Treacle) construct. DNA was stained with Hoechst 33342 (gray). Formation of Treacle condensates was induced with blue light illumination of the cells for 10 sec. Representative images are shown. **(E)** HeLa cells were transfected with Treacle-Katushka2S (Treacle-2S) at a quantity of 50 ng plasmids per 2×10^5^ cells. For low levels of expression analysis, the cells were fixed 16-24 hours after transfection. For high levels of expression analysis, the cells were fixed 48 hours after transfection. DNA was stained with DAPI (gray). Cells were analyzed by laser scanning confocal microscopy. Surface reconstructions of Treacle-2S FCs or extranucleolar condensates are shown in the right panels. Graphs illustrate quantification in arbitrary units of the 3D analysis of Treacle-2S condensate’s shape by LasX software. Sphere form factor = sphere area/particle area. **(F)** 16-24 h after transfection with Treacle-2S, HeLa cells were treated with 0.05µg/ml actinomycin D (AMD) to induce rDNA transcriptional repression and time-lapse imaged for 120 minutes. DNA was stained with Hoechst 33342 (gray). **(G)** 48 h after transfection with Treacle-2S, HeLa cells were live time-lapse imaged for 12 hours. DNA was stained with Hoechst 33342 (gray). Representative images of cells with Treacle-2S condensates fusion events are shown. Graphs illustrate the frequency of fusion Treacle condensates per cell for 12 hours. The calculation is presented for 6 biological replicates of 10 cells each. **(H)** HeLa cells were transiently transfected with Treacle-2S. 16-24 h after transfection half-FRAP analysis of Treacle-labelled fibrillar center (FCs) was performed. Half of the one FC was bleached, and fluorescence recovery was monitored. Representative time-lapse images of the photobleached FC are shown (magnified images). DNA was stained with Hoechst 33342 (gray). **(I)** Graphs illustrate the quantification of the FCs half-FRAP analysis described in (H). Half-FRAP curves represent the normalized intensity in the bleached half (blue) and the non-bleached half (green). Each trace represents an average of measurements for at least twenty FCs; error bars represent SD. **(J)** 16-24 h after transfection with Treacle-2S, HeLa cells were treated with 0.05µg/ml AMD to induce the formation of nucleolar caps. Half-FRAP analysis of Treacle-labelled nucleolar caps was performed. Representative time-lapse images of photobleached nucleolar caps are shown (magnified images). DNA was stained with Hoechst 33342 (gray). **(K)** Graphs illustrate the quantification of the nucleolar caps half-FRAP analysis described in (J). Half-FRAP curves represent the normalized intensity in the bleached half (blue) and the non-bleached half (green). Each trace represents an average of measurements for at least twenty caps; error bars represent SD. **(L)** HeLa cells were transiently transfected with Treacle-2S. 48 h after transfection, half-FRAP analysis of Treacle-2S extranucleolar condensates was performed. Half of one condensate was bleached, and fluorescence recovery was monitored. The analysis included condensates with diameters of up to 3 µm (type 1), as well as the largest condensates exceeding 5 µm in diameter (type 2). Representative time-lapse images of photobleached Treacle-2S extranucleolar condensates of both categories are shown (magnified images). DNA was stained with Hoechst 33342 (gray). **(M)** Graphs illustrate the quantification of the nucleolar caps half-FRAP analysis described in (L). Half-FRAP curves represent the normalized intensity in the bleached half (blue) and the non-bleached half (green). Each trace represents an average of measurements for at least twenty extranucleolar condensates of both categories; error bars represent SD. DNA was stained with Hoechst 33342 (gray).

To evaluate the ability of Treacle to self-assemble into biomolecular condensates, we first investigated its condensation properties using an *in vitro* system. For this purpose, a fragment of the Treacle protein (amino acids 291–426), encompassing two S/E-rich LCRs and two linker regions, was purified from Escherichia coli. *In vitro* experiments demonstrated that this recombinant fragment formed liquid-like condensates in the presence of 5% polyethylene glycol (Fig. 2B). These condensates exhibited a spherical morphology and frequently underwent fusion events (Fig. 2C).

To investigate Treacle’s ability to self-organize into biomolecular condensates within cells, we generated fusion proteins by linking Treacle to FusionRed (a non-self-dimerizing fluorescent protein) and *Arabidopsis thaliana* cryptochrome 2 (Cry2), which is known to oligomerize upon exposure to blue light, facilitating the formation of “optoDroplets” (Shin & Brangwynne, 2017). Expression of this opto–Treacle chimeric protein in HeLa cells revealed foci formation in nucleoli in the absence of blue light, consistent with Treacle’s natural tendency to occupy FCs (Fig. 2D; see Fig. S3B for the controls). Under blue light, opto–Treacle formed multiple small foci throughout the nucleoplasm (Fig. 2D).

Next, we explored Treacle’s ability to form biomolecular condensates when overexpressed as a fusion with the far-red fluorescent protein Katushka2S (referred to as 2S; Shcherbo et al., 2007) or green fluorescent protein (GFP). At low expression levels (16–24 hours post-transfection), Treacle’s fusion protein formed foci only in nucleoli, reflecting its natural occupation of FCs (Figs. 2E top panel, S3C). However, at increased expression levels (48–72 hours post-transfection), it began to form large spherical structures in the nucleoplasm (Figs. 2E bottom panel, S3D). The formation of these structures could not be attributed to the oligomerization of 2S or GFP, as comparable large nuclear condensates were observed at high levels of Treacle expression, even in the absence of a fused fluorescent protein (Fig. S3E). Both intranucleolar and extranucleolar Treacle–2S foci, induced by low and high levels of fusion protein expression respectively, exhibited a spherical shape with an aspect ratio close to one, suggesting susceptibility to surface tension (Fig. 2E). Transmission electron microscopy confirmed the internal homogeneity of Treacle condensates, showing circular structures with homogeneous protein content without any electron-dense or -light regions inside the condensates (Fig. S3F).

Like the *in vitro*-formed Treacle condensates, intracellular Treacle condensates demonstrated the ability to fuse. Our observations revealed that nucleolar caps form during AMD-induced (Figs. 2F, S3G; Movie S1) or DNA damage-induced (Fig. S3H; Movie S2) rDNA transcriptional repression due to the fusion of intranucleolar Treacle–2S foci. Prolonged live-cell imaging further demonstrated that extranucleolar biomolecular condensates from highly overexpressed Treacle–2S frequently fuse, indicating dynamic clustering (Fig. 2G; Movie S3).

Liquid condensates exhibit a high molecular exchange rate, often assessed through fluorescence recovery after photobleaching (FRAP; Ganser & Myong, 2020). To analyze the molecular dynamics of Treacle within its condensates, we employed a combination of full- and half-FRAP methods. Half-FRAP is a specific variation of partial FRAP, where half of the structure of interest is photobleached, unlike full-FRAP, which involves photobleaching the entire structure (Muzzopappa et al., 2022). Half-FRAP analysis revealed an increase in signal intensity in the bleached half of Treacle-2S-formed FCs, accompanied by a proportional decrease in fluorescence in the non-bleached half (Fig. 2H, I; Movie S4). Full-FRAP analysis of Treacle dynamics within FCs revealed a slow recovery rate (Fig. S4A), suggesting preferential internal mixing. This indicates that molecular exchange occurs between the two halves of the condensate without significant exchange across the boundary separating the condensate from the surrounding phase. Notably, fluorescence in the non-bleached region decreased to approximately half its initial value, suggesting near-complete internal mixing within the coacervate. Comparable full- and half-FRAP dynamics were observed for AMD-induced nucleolar caps (Figs. 2J, K, S4B; Movie S5) and extranucleolar condensates formed under elevated Treacle–2S expression (Figs. 2L, M, S4C; Movie S6). Furthermore, the largest extranucleolar condensates exhibited reduced Treacle molecular dynamics, suggesting more gel-like properties (Figs. 2L, M, S4D; Movie S7). Presumably, Treacle condensates undergo a liquid-to-gel phase transition over time or upon reaching a critical protein concentration within the condensate. Therefore, using diverse model systems, we illustrated that Treacle can generate biomolecular condensates with key features of structures formed through liquid-like phase separation.

### Treacle’s phase separation is regulated by its central and C-terminal domains

Since we have shown that an RD containing two S/E-rich LCRs can form liquid-like condensates *in vitro*, we next sought to investigate to what extent this condensation ability of RD is exerted intracellularly and to determine the roles of the CD and ND in intracellular Treacle condensation. For this purpose, we generated Treacle mutants missing the ND (Δ1–83), RD (Δ83–1121), or CD (Δ1121–1488) and overexpressed them in HeLa cells. We then assessed the condensation ability of the overexpressed deletion mutants in wild-type cells using several intracellular condensate models: the formation of (i) FCs and (ii) nucleolar caps under low expression levels (24 hours post-transfection) and (iii) large extranucleolar condensates under high expression levels (48 hours post-transfection).

The Δ1–83 mutant demonstrated condensation properties indistinguishable from full-length Treacle: it concentrated in FCs, formed stable nucleolar caps during transcriptional repression, and, upon increased expression levels, generated numerous large extranucleolar condensates (Figs. 3A, S5A, D, E). Using partial FRAP analysis, we further demonstrated that, similar to full-length Treacle (Fig. S5B, C), the Δ1–83 mutant exhibited a high rate of molecular exchange in all types of model condensates (Fig. 3B, C).

**Fig. 3.**
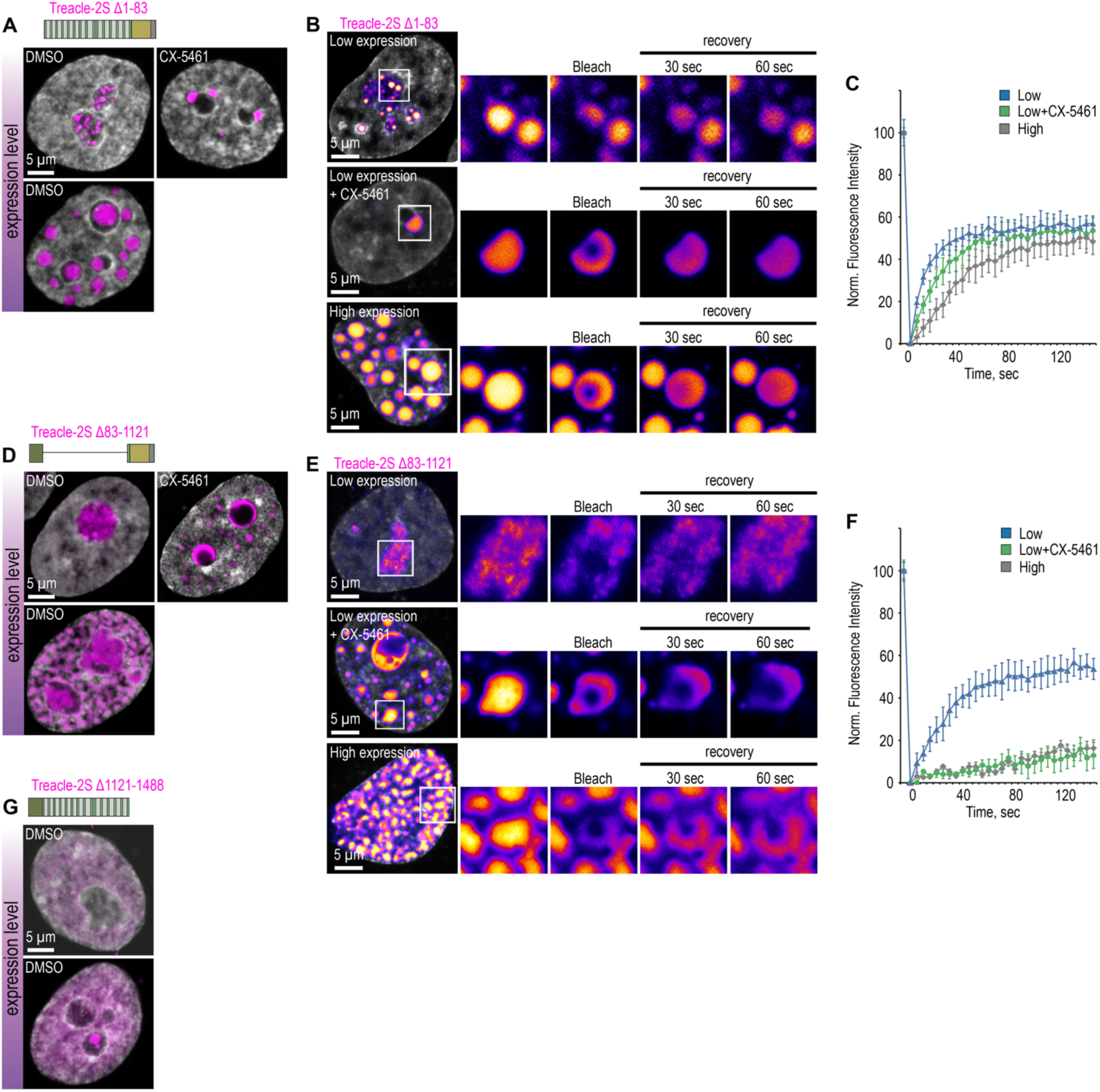
Treacle’s condensation is regulated by its central and C-terminal domains. **(A)** HeLa cells were transfected with Treacle-2S Δ1-83 deletion mutant. For low or high levels expression analysis cells were cultivated 16-24 or 48 h after transfection respectively. The expression level is indicated by the colored zone to the left of the cell images. HeLa cells with low expression level were additionally treated with CX-5461 to induce rDNA transcriptional repression and subsequent nucleolar cap formation. Cells were fixed and analyzed by laser scanning confocal microscopy. DNA was stained with DAPI (gray). Representative images of Treacle-2S Δ1-83 condensate are shown. **(B)** HeLa cells were transfected with Treacle-2S Δ1-83 deletion mutant and processed as described in (A). Partial FRAP analysis of Treacle-2S Δ1-83 condensates was performed. A part of each condensate type was photobleached, and the subsequent fluorescence recovery was monitored. Representative time-lapse images of the photobleached condensates are shown (magnified images). DNA was stained with Hoechst 33342 (gray). **(C)** HeLa cells were transfected with Treacle-2S Δ1-83 deletion mutant and processed as described in (A). Graphs illustrate the quantification of the Treacle-2S Δ1-83 condensates partial FRAP analysis described in (B). Each trace represents an average of measurements for at least twenty Treacle Δ1-83 condensates of each type; error bars represent SD. **(D)** HeLa cells were transfected with Treacle-2S Δ83-1121 deletion mutant and processed as described in (A). **(E)** HeLa cells were transfected with Treacle-2S Δ83-1121 deletion mutant and processed as described in (A). Partial FRAP analysis of Treacle-2S Δ83-1121 condensates dynamics was performed as described in (B). **(F)** HeLa cells were transfected with Treacle-2S Δ83-1121 deletion mutant and processed as described in (A). Partial FRAP analysis of 2S-fused Treacle Δ83-1121 condensates dynamics was performed as described in (B). Graphs illustrate the quantification of the Treacle-2S Δ83-1121 condensates partial FRAP analysis. Each trace represents an average of measurements for at least twenty Treacle-2S Δ83-1121 condensates of each type; error bars represent SD. **(G)** HeLa cells were transfected with Treacle-2S Δ1121-1488 deletion mutant fused with nuclear localization signal (NLS) from simian virus 40 (SV40). For low or high levels expression analysis cells were cultivated 16-24 or 48 h after transfection respectively. The expression level is indicated by the colored zone to the left of the cell images. Cells were fixed and analyzed by laser scanning confocal microscopy. DNA was stained with DAPI (gray). Representative images of cells are shown.

As expected, the Δ83–1121 mutant exhibited significantly compromised condensation properties. Under low expression levels, it failed to concentrate in FCs (Figs. 3D, S5F) and diffusely localized in the nucleolus while still maintaining a high molecular dynamics rate (Fig. 3E, F). During rDNA transcriptional repression, the Δ83–1121 mutant did not form classical nucleolar caps but partially redistributed between the nucleolar periphery and nucleoplasm, where it began to form condensates (Figs. 3D, S5F). Surprisingly, such relocation was associated with a change in its molecular dynamics state. Partial FRAP analysis revealed that nucleoplasmic condensates of the Δ83–1121 mutant, formed due to nucleolar transcriptional repression, began to transition into a solid-like state (Fig. 3E, F). A similar liquid-solid transition was observed for the Δ83–1121 mutant under high expression levels. In this case, it no longer formed distinct extranucleolar condensates but merged into a unified pan-nuclear solid network (Figs. 3D–F, S5F). This behavior suggests that the RD deletion alters Treacle’s interaction patterns, making its phase separation properties more reliant on rRNA and other nucleolar proteins.

Unexpectedly, the Δ1121–1488 mutant also failed to condense. When equipped with an artificial NLS from the SV40 virus, it successfully translocated into the nucleus; however, it remained diffusely distributed throughout the nucleoplasm, even under high expression conditions (Figs. 3G, S5G). Therefore, without the CD, Treacle cannot undergo autonomous condensation, likely relying on the CD as a nucleation factor. Collectively, these results suggest that the intracellular Treacle condensation depends on a cooperative interaction between RD and CD.

To gain further insights into these phenomena, we investigated the nature of interactions driving Treacle protein condensation. We assessed the behavior of Treacle intracellular condensates after treatment with 1,6-hexanediol, which disrupts hydrophobic interactions, or high salt concentrations, which interfere with electrostatic interactions. Treatment with the cell-permeable salt ammonium acetate disrupted both FCs and extranucleolar condensates formed at low and high Treacle expression levels, respectively (Fig. 4A). In contrast, treatment with 1,6-hexanediol did not affect the integrity of Treacle condensates (Fig. 4A). These observations were further validated using a condensation model based on the recombinant fragment of Treacle’s RD *in vitro*. The efficiency of Treacle’s fragment condensation *in vitro* remained unchanged in the presence of 1,6-hexanediol, whereas increasing salt concentration prevented condensation (Fig. 4B). Additionally, condensation of the Treacle RD fragment *in vitro* was sensitive to changes in pH. The optimal efficiency of condensation was observed at pH 7.0, whereas acidification of the medium decreased condensation, and alkalization resulted in the generation of insoluble aggregates (Fig. 4B). Therefore, it is clear that Treacle condensation, both *in vitro* and *in vivo*, is primarily governed by electrostatic interactions rather than hydrophobic forces.

**Fig. 4.**
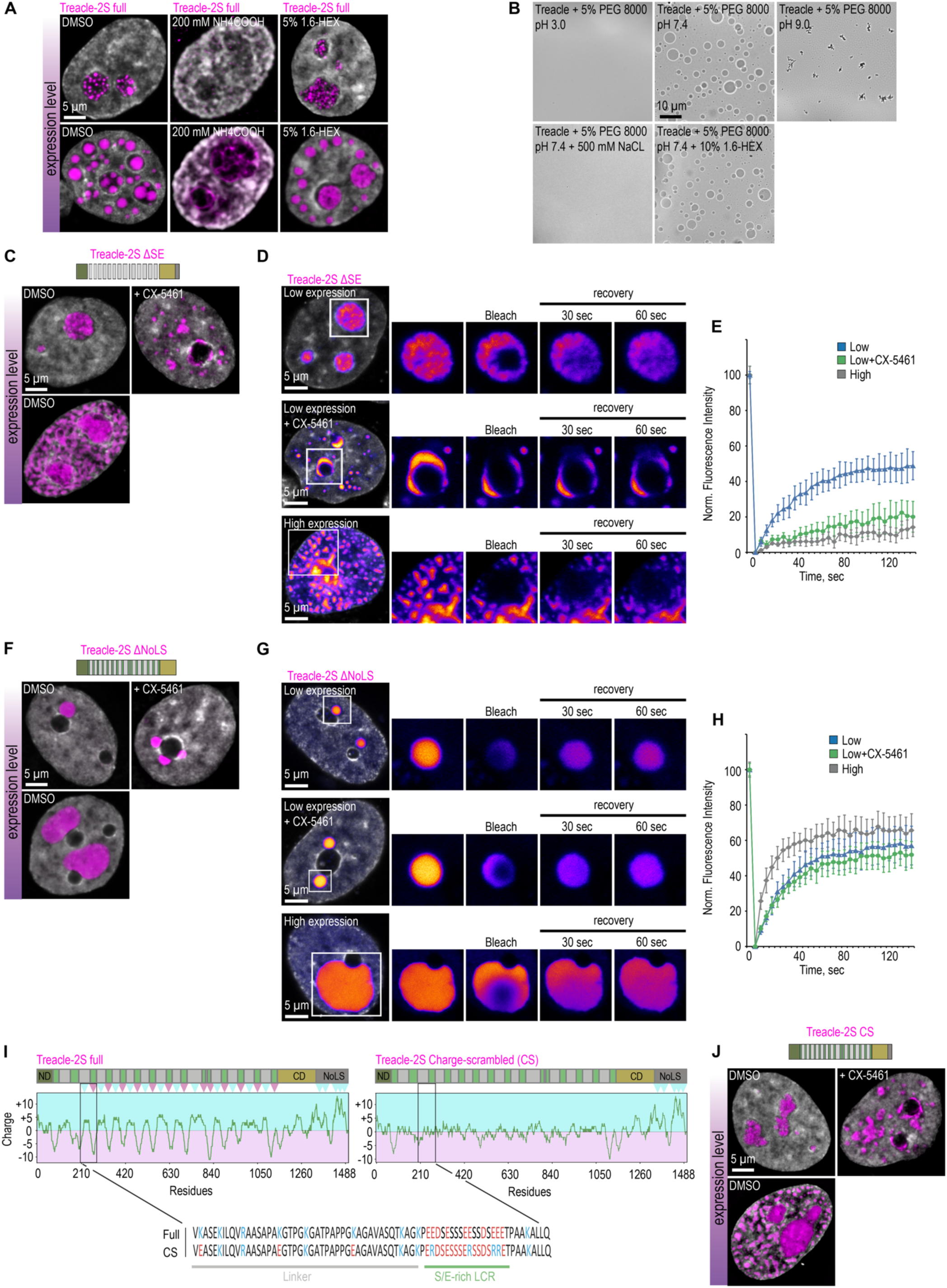
The condensation of Treacle is controlled by the specific charge distribution in its domains. **(A)** HeLa cells were transfected with Treacle-2S. For low or high levels expression analysis, cells were cultivated 16-24 or 48 h after transfection respectively. The expression level is indicated by the colored zone to the left of the cell images. Cells were treated with 5% 1,6-hexandiol (1,6-HEX) for 10 min or 200 mM ammonium acetate for 5 min. Cells were fixed and analyzed by laser scanning confocal microscopy. DNA was stained with DAPI (gray). Representative images of cells are shown. **(B)** The purified recombinant fragment of Treacle undergoes condensation *in vitro* in the presence of 10% 1,6-hexandiol (1.6-HEX), 500 mM sodium chloride or buffers with different pH. In each case, 5% PEG 8000 was used as a crowding agent. **(C)** HeLa cells were transfected with Treacle-2S ΔSE mutant. For low or high levels expression analysis cells were cultivated 16-24 or 48 h after transfection respectively. The expression level is indicated by the colored zone to the left of the cell images. HeLa cells with low expression level were additionally treated with CX-5461 to induce rDNA transcriptional repression. Cells were fixed and analyzed by laser scanning confocal microscopy. DNA was stained with DAPI (gray). Representative images of Treacle ΔSE condensate are shown. **(D)** HeLa cells were transfected with Treacle-2S ΔSE mutant and processed as described in (C). Partial FRAP analysis of Treacle-2S ΔSE condensates was performed. A part of each condensate type was photobleached, and the subsequent fluorescence recovery was monitored. Representative time-lapse images of the photobleached condensates are shown (magnified images). DNA was stained with Hoechst 33342 (gray). **(E)** HeLa cells were transfected with Treacle-2S ΔSE mutant and processed as described in (C). Graphs illustrate the quantification of the Treacle-2S ΔSE condensates partial FRAP analysis described in (D). Each trace represents an average of measurements for at least twenty Treacle-2S ΔSE condensates of each type; error bars represent SD. **(F)** HeLa cells were transfected with Treacle-2S Δ1350-1488 deletion mutant (Treacle-2S ΔNoLS) and processed as described in (C). **(G)** HeLa cells were transfected with Treacle-2S ΔNoLS deletion mutant and processed as described in (C). Partial FRAP analysis of Treacle-2S ΔNoLS condensates was performed as described in (D). **(H)** HeLa cells were transfected with Treacle-2S ΔNoLS deletion mutant and processed as described in (C). Partial FRAP analysis of Treacle-2S ΔNoLS condensates dynamics was performed as described in (D). Graphs illustrate the quantification of the Treacle-2S ΔNoLS condensates partial FRAP analysis. Each trace represents an average of measurements for at least twenty Treacle ΔNoLS condensates of each type; error bars represent SD. **(I)** Charge plots of full-length Treacle (left panel) and charge-scrambled Treacle (Treacle CS) form (right panel) are shown. Positive and negative charge blocks are depicted by blue and red triangles, respectively. Charge distribution was calculated as the sum of the charges (Arg and Lys, +1; Glu and Asp, −1;) in the 25 amino acids window range. The center of the panel shows the aligned amino acid sequences of one of the S/E-rich LCR and its adjacent linker of RD for both full Treacle and Treacle CS. **(J)** HeLa cells were transfected with Treacle-2S CS mutant and processed as described in (C).

It is reasonable to hypothesize that electrostatic interactions within Treacle condensates are facilitated by the presence of negatively charged S/E-rich LCRs in the RD and positively charged K-rich LCR in the CD. To evaluate this hypothesis, we generated a Treacle mutant lacking the 13 S/E-rich LCRs in the RD (ΔSE) and a Treacle mutant missing the 1350–1488 region (ΔNoLS), which includes the K-rich LCR. The deletion of the S/E-rich LCRs significantly compromised Treacle’s condensation properties. Like the Δ83–1121 mutant, Treacle ΔSE could not form FCs or nucleolar caps (Figs. 4C, S6A), remained in a liquid state within the nucleolus (Fig. 4D, E), and underwent a liquid-solid phase transition during nucleolar transcriptional repression or high overexpression levels (Fig. 4D, E). These findings indicate that the RD determines Treacle’s condensation properties through the S/E-rich LCRs. In turn, ΔNoLS mutant did not localize to the nucleolus but could form individual perinucleolar condensates (Figs. 4F, S6B). Interestingly, the volume of these condensates increased significantly with the expression level of ΔNoLS, but their number did not (Figs. 4F, S6B). This observation suggests that, in contrast to full-length Treacle, which forms a substantial number of extranucleolar condensates at high expression levels, ΔNoLS condensates exhibit a limited number of nucleation sites. This implies that the positively charged lysines in the K-rich LCR of the CD act as nucleation points for autonomous Treacle condensate formation. It is also noteworthy that all types of ΔNoLS condensates displayed high molecular dynamics (Fig. 4G, H). In contrast to full-length Treacle condensates, molecular exchange within ΔNoLS condensates occurred across the boundary separating the coacervate from the surrounding phase, rather than through internal mixing (Fig. S6C, D). This finding reinforces the role of the K-rich LCR as both a nucleation element in Treacle condensate formation and a regulator of Treacle’s volumetric dynamics within these condensates.

The disruption of Treacle’s condensation properties following the deletion of the S/E-rich LCRs underscores the significance of the negative charge within the central domain for phase separation. However, as noted earlier, the S/E-rich LCRs in the RD are not clustered but rather dispersed throughout the RD, interspersed with linker regions. We propose that for Treacle to exhibit proper condensation behavior, it is not merely the presence of a negative charge in the RD that is critical, both the presence of a negative charge in the RD and its specific spatial distribution within the molecule are critical.. Indeed, an analysis of the charge distribution along the Treacle amino acid sequence demonstrated that each S/E-rich LCR, in conjunction with its adjacent linker, within the RD constitutes a strong diblock ampholyte, where a positively charged block (linker) is followed by a negatively charged block (S/E-rich LCR; Fig. 4I left panel). In order to determine the physiological importance of this charge distribution, a variant of Treacle (Treacle CS) was created with the same overall net charge but scrambled blocks (Fig. 4I right panel). In Treacle CS, regions of opposite charge were removed while maintaining the same overall isoelectric point, amino acid composition, and positions of all other residues. As expected, scrambling the charges in the RD fully reproduced the condensation-defective phenotype observed in the Δ83–1121 and ΔSE mutants. Treacle CS failed to form FCs or nucleolar caps, instead assembling into a single pan-nuclear network at high overexpression levels (Fig. 4J, S6E). These findings suggest that Treacle’s condensation properties depend not only on the net negative charge within the RD but are critically influenced by the specific distribution of charges within this region.

In summary, our findings demonstrate that the cooperative interaction between the RD and CD governs the phase separation properties of Treacle, whereas the ND plays a negligible role in this process. The positively charged K-rich LCR within the CD facilitates its role as a nucleation site for Treacle condensate formation, while the alternating charge blocks in the RD stabilize Treacle condensates, ensuring that Treacle behavior remains independent of subnuclear localization or nucleolar transcriptional activity.

### Treacle condensation is essential for its proper interaction with nucleolar subcompartments

We next aimed to investigate the role of Treacle’s condensation properties in its functional interactions with the components of the FC, DFC, and GC within the nucleolus. First, we significantly downregulated endogenous Treacle expression (Treacle kd) using RNA interference (Fig. S7A) and then analyzed the structure organization of the FC, DFC, and GC within the nucleolus in the context of small interfering RNA (siRNA)-resistant full-length or condensation-defective (Δ83–1121 or CS) Treacle mutants.

The expressed full-length Treacle exhibited normal nucleolar localization, similar to endogenous Treacle. It co-localized within the FC with RPA194 and UBF, while FBL was localized in the DFC at the periphery of the FC, and NCL and B23 surrounded the DFC (Fig. 5A). In contrast, expression of the Δ83–1121 or CS Treacle mutants led to the dispersal and mixing of FC and DFC components within the nucleolus (Figs. 5B, S7B). Despite this, the Δ83–1121 or CS mutants still co-localized with UBF1, but to a significantly lesser extent with RPA194, suggesting a potential loss of their binding interaction. Interestingly, these mutants showed partial overlap with NCL, but not with B23 (Figs. 5B, S7B). These observations suggest that disrupting Treacle’s condensation properties may alter its interaction preferences..

**Fig. 5.**
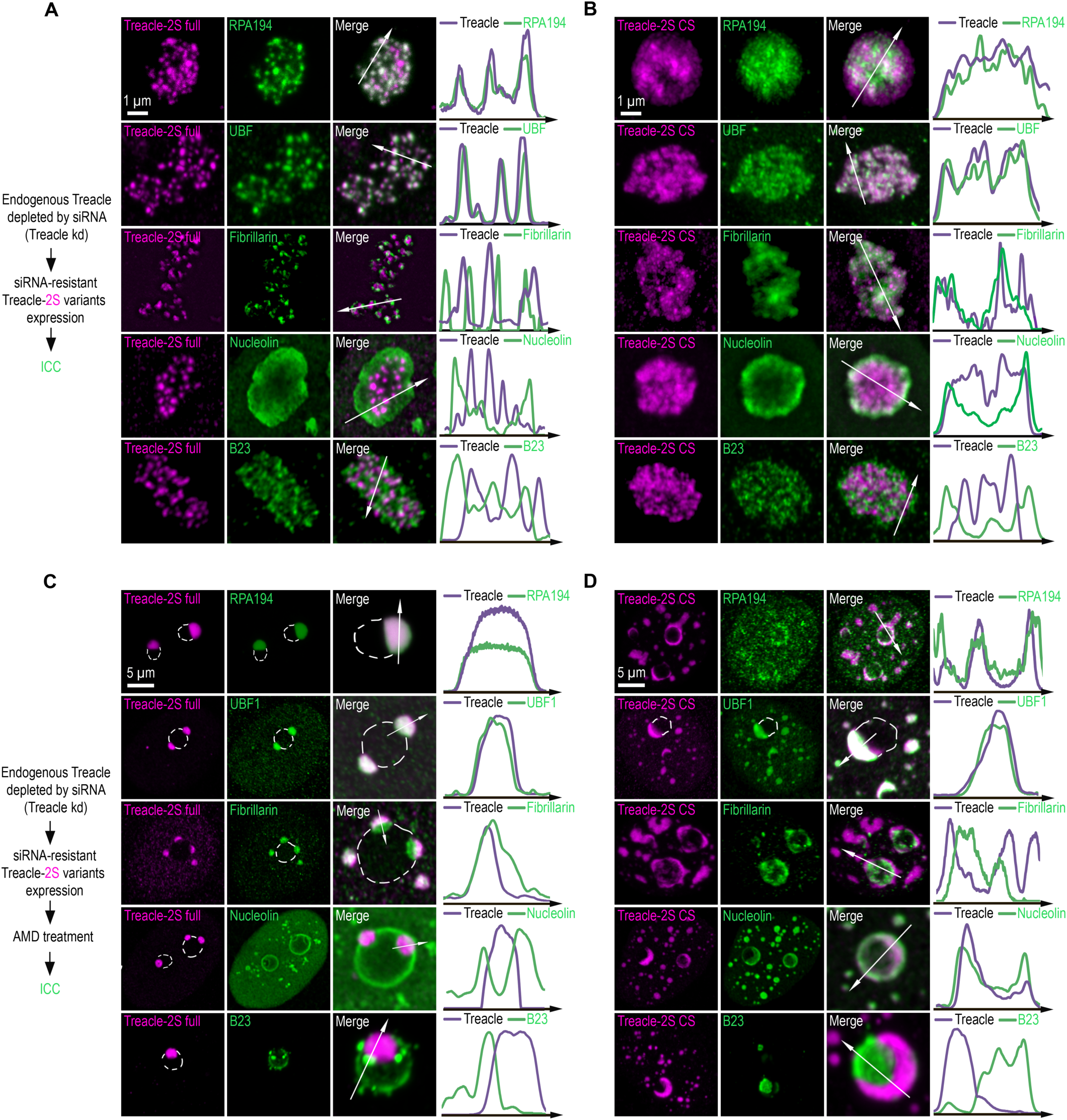
The condensation of Treacle facilitates its functional interactions with partner proteins. **(A)** Endogenous Treacle was depleted by siRNA-mediated knockdown (Treacle kd). Next, Treacle-depleted cells were transfected with siRNA-resistant full-length Treacle-2S (Treacle-2S full). Cells were fixed 16-24 after transfection and immunostained with either RPA194, UBF, Fibrillarin, B23 or Nucleolin antibodies. Cells were analyzed by laser scanning confocal microscopy. Representative images of magnified nucleoli are shown. Co-localization analysis was performed on the merged images. Graphs illustrate quantification in arbitrary units of Treacle-2S variants and RPA194, UBF, Fibrillarin, B23 or Nucleolin fluorescence distribution along the lines shown in the figures. **(B)** Endogenous Treacle was depleted by siRNA-mediated knockdown (Treacle kd). Next, Treacle-depleted cells were transfected with siRNA-resistant charge-scrambled Treacle-2S (Treacle-2S CS) and processed as described in (A). **(C)** Endogenous Treacle was depleted by siRNA-mediated knockdown (Treacle kd). Next, Treacle-depleted cells were transfected with siRNA-resistant full-length Treacle-2S (Treacle-2S full). 16-24 h after transfection cells were treated with 0.05µg/ml actinomycin D (AMD) to induce rDNA transcriptional repression, fixed and processed as described in (A). **(D)** Endogenous Treacle was depleted by siRNA-mediated knockdown (Treacle kd). Next, Treacle-depleted cells were transfected with siRNA-resistant charge-scrambled Treacle-2S (Treacle-2S CS). 16-24 h after transfection cells were treated with 0.05µg/ml actinomycin D (AMD) to induce rDNA transcriptional repression, fixed and processed as described in (A).

This hypothesis is further supported by the analysis of the localization of nucleolar components and condensation-defective Treacle forms following AMD-induced nucleolar transcriptional repression. After AMD treatment, full-length Treacle co-localized with UBF1 and RPA194 at the nucleolar caps (Fig. 5C). FBL partially merged with the caps, while NCL and B23 were completely displaced to the nucleolar rim and nucleoplasm (Fig. 5C). In contrast, the AMD-induced perinucleolar or nucleoplasmic solid condensates formed by the Δ83–1121 or CS Treacle mutants still co-localized with UBF1, but to a significantly lesser extent with RPA194 (Figs. 5D, S7C). Interestingly, these condensates exhibited behavior opposite to that of full-length Treacle condensates with respect to FBL and NCL: they displaced FBL and incorporated NCL (Figs. 5D, S7C). The patterns of association between FC/DFC/GC proteins and the extranucleolar condensates of Δ83–1121 and CS Treacle mutants at high expression levels exhibited similar alterations when compared to those observed with full-length Treacle (Fig. S7D, E).

Thus, the disruption of Treacle’s condensation properties, either through RD deletion or charge scrambling, results in a shift in its interaction repertoire. Specifically, while interaction with UBF1 is preserved, interactions with RNA Pol I and FBL are notably diminished, whereas a new interaction with NCL is established. It is likely that the correct condensation of Treacle contributes to the formation of the cooperative FC/DFC structure by mediating functional interactions with partner proteins.

### The condensation of Treacle regulates rRNA transcription and processing

In the above sections, we demonstrated that Treacle supports both rRNA transcription and processing. We hypothesized that this multifunctionality of Treacle could be related to its ability to condense and the formation of the cooperative FC/DFC structure. To test this hypothesis, we significantly downregulated endogenous Treacle expression (Treacle kd) using RNA interference and then analyzed rRNA transcription levels in the context of siRNA-resistant full-length or condensation-defective (Δ83– 1121 or CS) Treacle mutants. We found that the expression of condensation-defective Treacle mutants significantly reduced the transcription of rRNA compared to full-length Treacle (Fig. 6A). This effect correlated with a reduced ability of condensation-defective Treacle mutants to maintain the association of partner proteins RPA194 and UBF1 at the rDNA promoter (Fig. 6C), along with diminished intrinsic binding of these mutants to rDNA (Fig. 6B).

**Fig. 6.**
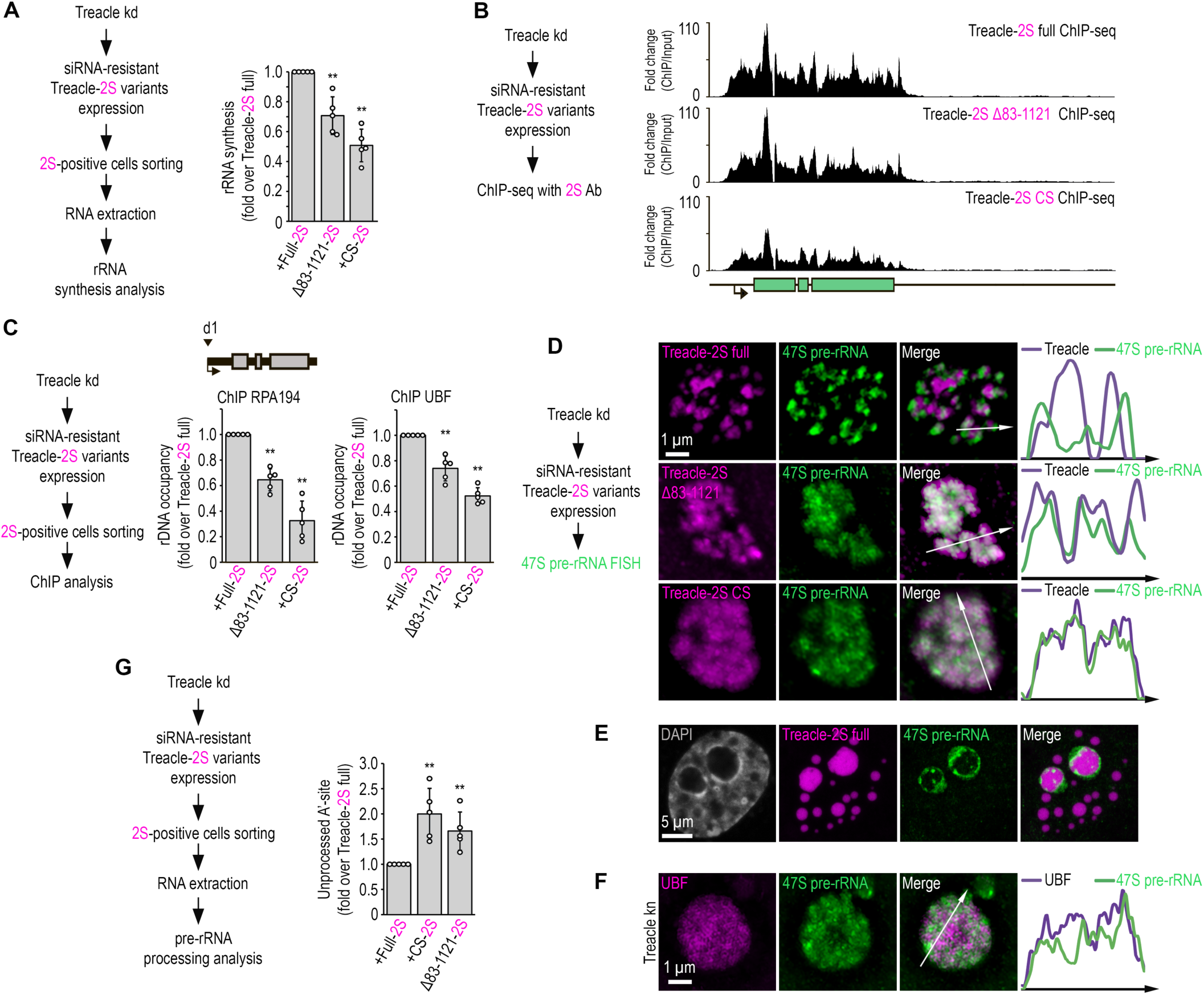
Treacle LLPS is essential for the transcription and processing of rRNA. **(A)** Endogenous Treacle was depleted by siRNA-mediated knockdown (Treacle kd). Next, Treacle-depleted cells were transfected with siRNA-resistant full-length Treacle-2S (Treacle-2S full), charge-scrambled Treacle-2S (Treacle-2S CS) or Treacle-2S Δ83-1121 deletion mutant (Treacle-2S Δ83-1121). Cells were fixed 16-24 after transfection and subjected to cell sorting in the fluorescent analysis mode to obtain 2S-positive populations. The sorted cell fractions were used for RNA extraction. RT-qPCR was performed; it shows levels of 47S pre-rRNA normalized to GAPDH mRNA. Normalized pre-rRNA level in full-length Treacle-2S-positive cells is set to 1. Values are mean ± SD. The calculation is presented for 5 biological replicates. **, p < 0.01 by unpaired t test. **(B)** HeLa cells were processed as described in (A). Cells were fixed 16-24 after transfection and subjected to ChIP-seq analysis with Katushka2S antibodies. ChIP-seq signal were normalized to the input. **(C)** HeLa cells were processed as described in (A). Cells were fixed 16-24 after transfection and subjected to cell sorting in the fluorescent analysis mode to obtain 2S-positive populations. The sorted cell fractions were used for used for ChIP with RPA194 or UBF antibodies. ChIP was followed by qPCR using the d1 primers to the promoter of the rRNA gene (positioned as indicated on the scheme). Percentage of enrichment relative to input in full-length Treacle-2S-positive cells is set to 1.Values are mean ± SD. The calculation is presented for 5 biological replicates. **, p < 0.01 by unpaired t test. **(D)** HeLa cells were processed as described in (A). Cells were fixed 16-24 after transfection and stained for 47S pre-rRNA (revealed by single-molecule FISH, smFISH). Cells were analyzed by laser scanning confocal microscopy. Representative images of magnified nucleoli are shown. Co-localization analysis was performed on the merged images. Graphs illustrate quantification in arbitrary units of Treacle-2S variants and smFISH fluorescence distribution along the lines shown in the figures. **(E)** HeLa cells were transfected with full-length Treacle-2S. 48 h after transfection cells were fixed and stained for 47S pre-rRNA (revealed by single-molecule FISH, smFISH). Cells were analyzed by laser scanning confocal microscopy. Representative images of magnified nucleoli are shown. **(F)** HeLa cells were transfected with a construct coding CRISPR/Cas9 and sgRNA to the TCOF1 gene (Treacle kn). After 7-10 days after transfection, the cells were fixed and stained with UBF antibodies and 47S pre-rRNA. Representative images of Treacle-negative nucleolus (magnified images) are shown. **(G)** HeLa cells were processed as described in (A). Cells were fixed 16-24 after transfection and subjected to cell sorting in the fluorescent analysis mode to obtain 2S-positive populations. The sorted cell fractions were used for RNA extraction. RT-qPCR was performed; it shows levels of A’-site contained unprocessed rRNA normalized to GAPDH mRNA. Normalized unprocessed rRNA level in full-length Treacle-2S-positive cells is set to 1. Values are mean ± SD. The calculation is presented for 5 biological replicates. **, p < 0.01 by unpaired t test.

These observations suggest that the condensation properties of Treacle underlie its ability to concentrate components of the transcriptional machinery in the FC and separate them from the DFC. Moreover, such separation is likely required to maintain highly efficient transcription of rDNA. We hypothesized that the same principle could regulate the role of Treacle in rRNA processing. To test this assumption, we depleted the level of endogenous Treacle using RNA interference and then analyzed the spatial localization of newly synthesized 47S pre-rRNA and its processing level in cells expressing siRNA-resistant full-length or condensation-defective (Δ83–1121 or CS) Treacle mutants. Microscopy analyses with fluorescence *in situ* hybridization (FISH)-labeled rRNA revealed that newly synthesized 47S pre-rRNA transcripts clustered at the periphery of FCs in full-length Treacle-expressing cells, reflecting the radial flow of rRNA from the FC to the GC (Fig. 6D). Newly synthesized rRNA was also relocated to the periphery of large Treacle condensates formed at high expression levels (Fig. 6E). However, no apparent clustering of newly synthesized rRNA was observed in cells expressing condensation-defective Treacle mutants. Instead, the newly synthesized transcripts were diffusely mixed with Δ83–1121 and CS Treacle (Fig. 6D). Nascent rRNA transcripts were similarly mixed with delocalized UBF in cells depleted of endogenous Treacle (Treacle kn; Fig. 6F). Finally, reverse transcription-quantitative polymerase chain reaction (RT-qPCR) analysis confirmed that cells overexpressing condensation-defective Treacle mutants had impaired 5′ external transcribed spacer processing compared to cells overexpressing full-length Treacle (Fig. 6G). Therefore, it can be concluded that the mixing of FC and DFC components due to the expression of condensation-defective Treacle or the depletion of endogenous Treacle disrupts the directional traffic of nascent rRNA from the FC to the DFC and further to the GC, potentially causing inefficient processing.

Our results indicate that Treacle’s condensation properties separate the FC and DFC, leading to the spatial segregation of rRNA synthesis and subsequent processing. The mixing of FC and DFC components due to the disruption of Treacle’s condensation ability reduces the efficiency of both processes, with equivalent effects to the complete depletion of endogenous Treacle.

### The condensation of Treacle is essential for DDR activation in ribosomal genes under genotoxic stress

Previous studies have demonstrated that Treacle is critical for inducing the DDR in ribosomal genes under certain types of stress (Korsholm et al., 2019; Larsen et al., 2014; Mooser et al., 2020; Velichko et al., 2019, 2021). Here, we aimed to investigate the contribution of Treacle’s condensation to rDNA damage response. We induced rDNA damage using the widely used chemotherapeutic drug etoposide (VP16), which acts as a topoisomerase II inhibitor, inducing double-strand breaks (DSBs) (Bax et al., 2019). Treating cells with 90 µM VP16 for 30 minutes resulted in the rapid recruitment of TOPBP1 to nucleoli and its colocalization with Treacle in HeLa (Figs. 7A, S8A) and MCF7 cells (Fig. S8B). Furthermore, proximity ligation assay (PLA) analysis conducted after DNA damage induction provided evidence of a physical association between TOPBP1 and Treacle (Figs. 7B, S8C). Reducing endogenous Treacle levels, either through RNA interference (Treacle kd) or CRISPR/Cas9-mediated depletion (Treacle kn), effectively blocked the relocalization of TOPBP1 to nucleoli (Figs. 7D, S9A–B) and its occupancy of rDNA (Fig. 7C). Interestingly, with Treacle kd, a substantial amount of Treacle still remained in nucleoli (Fig. 7D). However, this amount was insufficient to facilitate its interaction with TOPBP1, likely due to the need for specific stoichiometric ratios for efficient interaction.

**Fig. 7.**
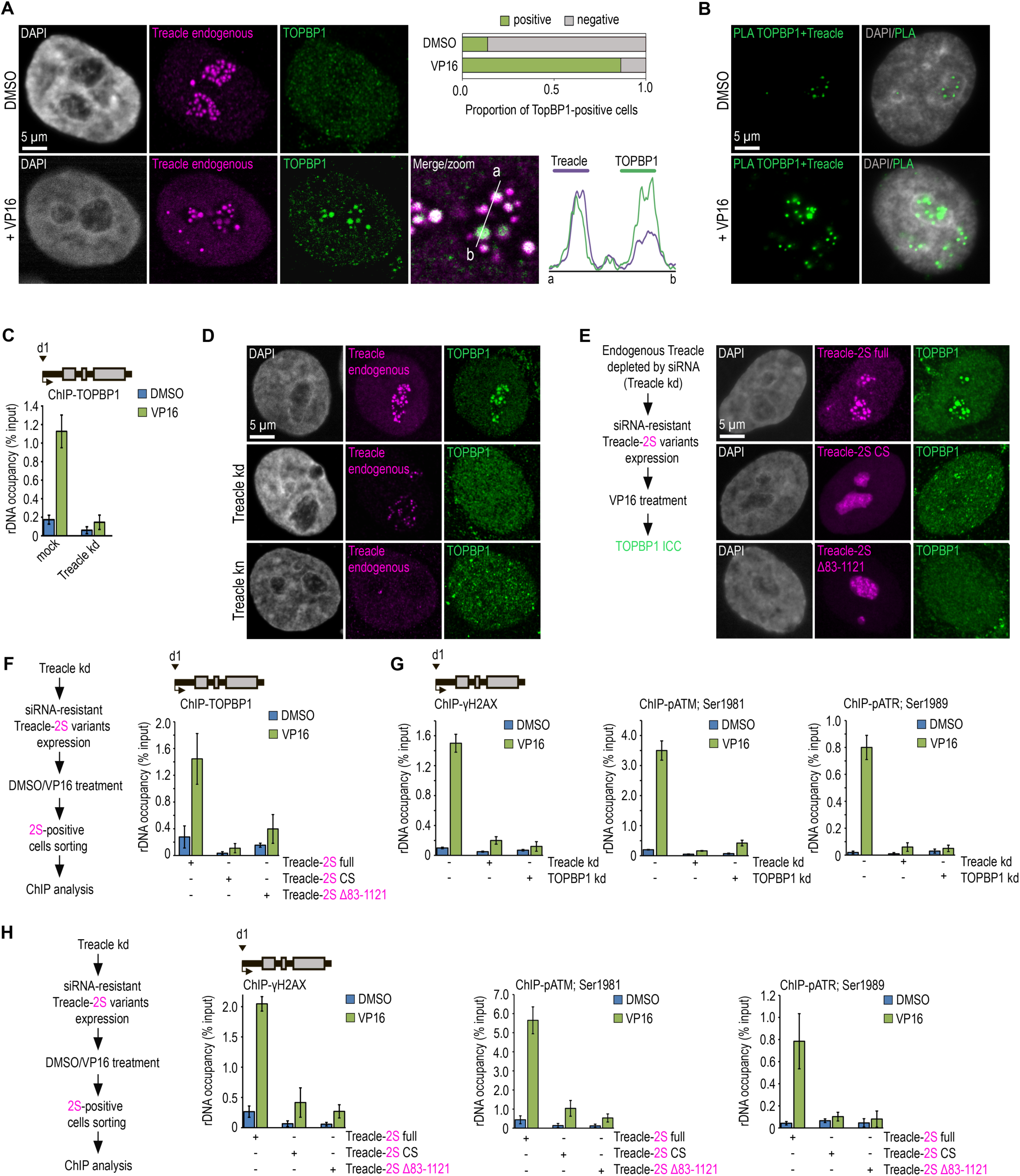
Treacle phase separation is essential for DDR activation in ribosomal genes under genotoxic stress conditions. **(A)** DMSO-treated and VP16-treated (90 μM, 30 min) HeLa cells were co-immunostained for Treacle (Treacle endogenous; magenta) and TOPBP1 (green) and analyzed by laser scanning confocal microscopy. The DNA was stained with DAPI (gray). Co-localization analysis was performed on the merged images (magnified images). Graphs illustrate quantification in arbitrary units of Treacle and TOPBP1 fluorescence distribution along the lines shown in the figures. Percentage of cells containing TOPBP1 (TOPBP1-positive) foci within nucleoli is shown. **(B)** DMSO-treated and VP16-treated (90 μM, 30 min) HeLa cells were subjected to Proximity Ligation Assay (PLA) with antibodies against TOPBP1 and Treacle. DNA was stained with DAPI (gray). PLA detection of Treacle-TOPBP1 interactions is visible as distinct green fluorescent dots. **(C)** Mock-treated HeLa cells or cells with siRNA-mediated Treacle knockdown (Treacle kd) were treated with DMSO or VP16 (90 μM, 30 min). ChIP experiments were performed with antibodies against TOPBP1. ChIP was followed by qPCR using the d1 primers to the promoter of the rRNA gene (positioned as indicated on the scheme). Data are represented relative to the input. Values are means ±SD from at least three independent replicates. **(D)** Intact HeLa cells, siRNA-depleted for Treacle (Treacle kd) cells or CRISPR/Cas9-depleted for Treacle (Treacle kn) cells were treated with 90 μM VP16 for 30 min. Cells were co-immunostained for TOPBP1 (green) and Treacle (magenta) antibodies and analyzed by laser scanning confocal microscopy. The DNA was stained with DAPI (gray). **(E)** Endogenous Treacle was depleted by siRNA-mediated knockdown (Treacle kd). Next, Treacle-depleted cells were transfected with plasmid constructs encoding either siRNA-resistant full-length Treacle-2S (Treacle-2S full), Treacle-2S Δ83-1121 deletion mutant, or charge-scrambled mutant Treacle-2S (Treacle-2S CS). 24 h after transfection, cells were treated with VP16 (90 μM for 30 min), fixed and stained for TOPBP1 (green) and analyzed by laser scanning confocal microscopy. The DNA was stained with DAPI (gray). **(F)** Endogenous Treacle was depleted by siRNA-mediated knockdown (Treacle kd). Treacle-depleted cells were transfected with plasmid constructs encoding either siRNA-resistant Treacle-2S full, Treacle-2S Δ83-1121, or Treacle-2S CS. 24 h after transfection, cells were treated with DMSO or VP16 (90 μM for 30 min) and fixed. Cells were subjected to cell sorting in the fluorescent analysis mode to obtain 2S-positive populations. At least 2×10^6^ sorted cells were used for ChIP with TOPBP1 antibodies. ChIP was followed by qPCR using the d1 primers to the promoter of the rRNA gene (positioned as indicated on the scheme). Data are represented relative to the input. Values are means ±SD from at least three independent replicates. **(G)** Intact HeLa cells and cells siRNA-depleted for either Treacle (Treacle kd) or TOPBP1 (TOPBP1 kd) were treated with DMSO or 90 μM VP16 for 30 min. ChIP experiments were performed with antibodies against phospho-ATR (pATR; Ser1989), phospho-ATM (pATM; Ser1981) or γH2AX antibodies. ChIP was followed as described in (F). **(H)** HeLa cells were processed as described in (F). At least 2×10^6^ sorted cells were used for ChIP with phospho-ATR (pATR; Ser1989), phospho-ATM (pATM; Ser1981) or γH2AX antibodies. ChIP was followed by qPCR using the d1 primers to the promoter of the rRNA gene (positioned as indicated on the scheme). Data are represented relative to the input. Values are means ±SD from at least three independent replicates.

Next, we investigated whether the interaction between Treacle and TOPBP1 in Treacle kd cells could be restored by overexpressing different Treacle variants. Immunocytochemical analysis and chromatin immunoprecipitation with antibodies against TOPBP1 indicated that expressing (siRNA)-resistant full-length Treacle in Treacle kd cells effectively restored the VP16-induced interaction between Treacle and TOPBP1 (Figs. 7E, S10A), as well as the enrichment of TOPBP1 at the promoters of rDNA (Fig. 7F). However, expressing (siRNA)-resistant condensation-defective forms of Treacle (Δ83–1121 or CS) did not lead to such restoration (Figs. 7E–F, S10A).

The response to DNA damage involves the activation of various signaling kinases and the recruitment of different repair factors to the DNA break site. Chromatin immunoprecipitation followed by qPCR with antibodies against a panel of DNA repair proteins confirmed that VP16 induced DDR at the promoters of rDNA, including the activation of ATM serine/threonine kinase (ATM) and ATR, phosphorylation of the H2A.X variant histone (H2AX), and recruitment of repair factors such as tumor protein p53 binding protein 1 (TP53BP1/53BP1) and BRCA1 DNA repair associated (BRCA1; Figs. 7G, S10C). As expected, nucleolar DDR was fully controlled by the interaction between Treacle and TOPBP1, as the knockdown of either of these proteins dramatically reduced VP16-induced repair signaling at rDNA (Figs. 7G, S10B–C). Finally, it was confirmed that nucleolar DDR could be efficiently restored in Treacle kd cells by overexpressing (siRNA)-resistant full-length Treacle but not its condensation-defective mutants (Δ83–1121 or CS; Figs. 7H, S10D). Therefore, we can conclude that Treacle’s phase separation is crucial in facilitating its binding to TOPBP1 and activating the DDR in response to genotoxic damage to rDNA.

## Discussion

The nucleolus is a multicomponent phase condensate formed on the platform of rRNA, which is processed and incorporated into assembling ribosomes simultaneously with migration from FCs to DFCs and further to GCs. The constitutive components of DFCs and GCs are NCL and NPM1, respectively (Feric et al., 2016; Lafontaine et al., 2021). Which protein plays a comparable role in FCs continues to be debated. Our results suggest that the phosphoprotein Treacle serves as the scaffold protein for nucleolar FCs in human cells. This conclusion aligns entirely with the observation that the evolutionary emergence of Treacle correlates with the appearance of the three-partite nucleolus in amniotes (Jaberi-Lashkari et al., 2023). Introducing human Treacle into zebrafish, which lack a competent ortholog for FC formation, induced the reorganization of the nucleolar structure from bipartite to tripartite, accompanied by the formation of FC-like structures.

Our findings demonstrate that the condensation properties of Treacle are governed by the synergistic interplay between its RD and CD. While the RD can form liquid-like condensates alone *in vitro*, its condensation within the cellular environment requires the positively charged CD, which likely functions as a nucleation platform. Notably, the zebrafish ortholog of Treacle exhibits sequence homology with human Treacle exclusively within the RD (Hill-Terán et al., 2024), lacking a homologous sequence in the CD. This structural difference likely accounts for its inability to undergo condensation and form FC-like structures.

Treacle functions as a block polyampholyte, with its condensation properties governed by the alternating charges of the CD and the strongly positive charge of the C-terminal domain. These properties suggest that its condensation is likely driven by coacervation or self-coacervation mechanisms. Complex coacervation refers to a phase transition process mediated by polyelectrolyte complexation, where oppositely charged macromolecules associate through electrostatic interactions and phase-separate from the surrounding solution to form dense, charge-balanced structures known as coacervates (Choi et al., 2020; Sathyavageeswaran et al., 2024; Sing & Perry, 2020; Pappu et al., 2023). The driving forces for complex coacervation can stem from homotypic or heterotypic interactions between charged polymers (Sanders et al., 2020). Our experimental findings suggest that both mechanisms may be drivers of Treacle’s coacervation.

Homotypic coacervation of Treacle may occur through a two-step process: nucleation is facilitated by electrostatic interactions between the lysine (K)-rich low-complexity region (LCR) in the CD and the glutamic acid (E)-rich LCR in the RD, while fluidization is achieved via dynamic interactions between the E-rich LCR and K-rich linkers, primarily within the RD. Conversely, heterotypic coacervation of Treacle may involve interactions with other polyampholyte proteins enriched in the FCs, many of which are known Treacle partners and possess the ability to facilitate coacervation. For example, it is hypothesized that the CD of Treacle may promote nucleation through interactions with UBF, which contains extended aspartic acid (D)/glutamic acid (E) tracts and demonstrates autonomous condensation properties in the presence of rDNA (Gal et al., 2022; King et al., 2024; Lin & Yeh, 2009). Additionally, Treacle interacts with RNA Pol I, whose subunits (e.g., POLR1F and POLR1G) are characterized by prominent K-rich regions and E-rich tracts (King et al., 2024). These regions are likely to facilitate electrostatic interactions between RNA Pol I and Treacle, potentially modulating its condensation dynamics within the cellular environment. This hypothesis is further supported by our experimental observations showing that charge scrambling or deletion of Treacle’s RD partially disrupts its interaction with RNA Pol I subunit RPA194, reduces its recruitment to rDNA, and impairs its condensation properties. Together, these findings highlight the interplay of homotypic and heterotypic interactions in driving Treacle’s coacervation and its functional role within the nucleolus.

Endogenous Treacle may undergo post-translational modifications. Specifically, the RD of Treacle is known to be heavily phosphorylated and ubiquitinated (Mooser et al., 2020; Werner et al., 2015, 2018). Currently, the role of these modifications is primarily attributed to facilitating interactions with partner proteins such as the cullin 3 (CUL3), kelch repeat and BTB domain containing 8 (KBTBD8) complex or nucleolar and coiled-body phosphoprotein 1 (NOLC1; Werner et al., 2018). However, it is known that phosphorylation can modulate phase separation, particularly by enhancing or reducing the polyampholytic properties of disordered regions (Yamazaki et al., 2022). We propose that post-translational modifications, such as ubiquitination and phosphorylation, fine-tune the condensation properties of Treacle. These modifications may do so by modulating interactions with binding partners and altering the ampholytic properties of the RD through changes in its charge ratio.

The 47S pre-rRNA is synthesized at the FC/DFC boundary, while its primary processing begins in the DFC. DFC formation is mediated by the phase separation of FBL gathered on the rRNA scaffold and is maintained as long as there is a directed radial flow of rRNA from the FC to the DFC (Wu et al., 2021; Yao et al., 2019). While the role of Treacle in ensuring rRNA processing has been repeatedly described (Calo et al., 2018; Gonzales et al., 2005), the underlying mechanism remains unclear. We demonstrated that the efficiency of rRNA processing is compromised not only by the complete depletion of endogenous Treacle but also by the expression of condensation-defective Treacle mutants. This impairment correlates with a loss of Treacle association with fibrillarin and, conversely, an increased association of Treacle with newly synthesized rRNA and nucleolin. This shift likely disrupts the cooperative organization of the FC/DFC and the mixing of their components. The resulting disorganization and mixing of FC and DFC components, driven by the loss of Treacle’s condensation properties, may compromise the radial flow of rRNA, thereby hindering its normal processing.

We also explored the role of Treacle’s phase separation in nucleolar DDR. Treacle’s well-known partner, TOPBP1, plays a crucial role in activating ATR during DNA replication stress (Day et al., 2021). The interaction between TOPBP1 and Treacle is essential for nucleoli-specific DDR and replication stress response (Mooser et al., 2020; Velichko et al., 2019, 2021). This interaction occurs via phosphoserines 1227 and 1228 in a conserved acidic motif in the CD of Treacle (Mooser et al., 2020). However, we showed that Treacle with intact phosphoserines 1227/1228 but unable to undergo condensation loses the ability to interact with TOPBP1 during genotoxic stress. This observation implies that Treacle’s interaction with TOPBP1 and the subsequent activation of nucleolar DDR requires Treacle’s phase separation ability.

In conclusion, our study revealed Treacle’s role not only as a structural scaffold for the FC/DFC but also as a nucleolar hub protein that integrates functions in rDNA transcription, rRNA processing, and rDNA repair.

## Materials and methods

### Cell culture and drug treatment

Human HeLa (ATCC® CCL-2™), MCF7 (ATCC® HTB-22) and human skin fibroblasts (female 46XX) were cultured in DMEM (PanEco) supplemented with 10% fetal bovine serum (FBS; HyClone/GE Healthcare) and penicillin/streptomycin. The cells were cultured at 37°C in a conventional humidified CO_2_ incubator. DNA damage was induced by the treatment of cells with 90 µM etoposide (Sigma-Aldrich, #E1383) for 30 min or 1 h. For rRNA transcription inhibition, cells were treated with 0,05 µg/ml actinomycin D (Sigma-Aldrich, #A1410) or 20 µM CX-5461 for 3 h. To obtain 1,6-hexanediol (HD)-treated cells, HeLa cells were incubated with 5% 1,6-HD (Sigma-Aldrich, #240117) in serum-free medium at 37°C in a humidified atmosphere for 10 min. To obtain ammonium acetate-treated cells, HeLa cells were incubated with 200 mM ammonium acetate in a complete culture medium at room temperature for 5 min.

### Plasmid constructs

For the FusionRED-Treacle-Cry2 (opto-Treacle) construct, the full-length Treacle was amplified by PCR from cDNA with primer set #1 (Supplementary Table 1) using KAPA High-Fidelity DNA Polymerase (KAPA Biosystems, KE2502). The forward and reverse primers contained XhoI sites. The amplified fragment was inserted into the FusionRed-C vector (Evrogen, FP411) linearized with XhoI. The Cry2 fragment was amplified by PCR from the plasmid pHR-mCh-Cry2WT (Addgene, #101221) with primer set #2 (Supplementary Table 1) digested with NheI and inserted into pFusionRed-Treacle linearized with NheI using NheI enzyme.

For the FusionRED-FUS-Cry2 construct, the FUS was amplified by PCR from the plasmid pHR-FUSN-mCh-Cry2WT (Addgene, #101223) with primer set #3 (Supplementary Table 1) using KAPA High-Fidelity DNA Polymerase (KAPA Biosystems, KE2502). The forward and reverse primers contained BspEI and KpnII sites respectively. The amplified fragment was inserted into the pFusionRed-C vector (Evrogen, FP411). The Cry2 fragment was amplified by PCR from the plasmid pHR-mCh-Cry2WT (Addgene, #101221) with primer set #2 (Supplementary Table 1) digested with NheI and inserted into pFusionRed-Treacle using NheI enzyme.

To generate Treacle-GFP or Treacle-Katushka2S constructs, the full-length Treacle was amplified by PCR from cDNA with primer set #4 (Supplementary Table 1) using KAPA High-Fidelity DNA Polymerase (KAPA Biosystems, KE2502). The forward and reverse primers contained BglII and BamHI sites respectively. The amplified fragment was inserted into the pTurboGFP-C (Evrogen, FP511) or pKatushka2S-C (Evrogen, FP761) vectors using BglII/BamHI restriction/ligation.

Treacle Δ1-83 deletion mutant was constructed based on the pTreacle-GFP full or pTreacle-2S full plasmid using iProof High-Fidelity DNA polymerase with primer set #5 (Supplementary Table 1). The resulting DNA template after PCR was reprecipitated and treated with the DpnI restriction enzyme. In the next step, the desired DNA template was purified on an agarose gel, phosphorylated with T4 Polynucleotide Kinase (T4 PNK), and ligated.

To obtain the charge-scrambled Treacle-GFP or Treacle-2S (Treacle-GFP CS or Treacle-2S CS), a fragment encoding 137-1130 aa of Treacle was removed from the pTreacle-GFP or pTreacle-2S plasmid using EcoR1 and HindIII restrictases. This fragment was then replaced with a synthetic sequence encoding the amino acid sequence with the necessary changes (see Supplementary Table 2).

To obtain the Treacle ΔSE mutant, a fragment encoding 137-1130 aa of Treacle was removed from the pTreacle-GFP or pTreacle-2S plasmid using EcoR1 and HindIII restrictases. This fragment was then replaced with a synthetic sequence encoding the amino acid sequence with the necessary changes (see Supplementary Table 2).

Treacle Δ83-1121 deletion mutant was constructed based on the pTreacle-GFP full or pTreacle-2S full plasmid using iProof High-Fidelity DNA polymerase with primer set #6 (Supplementary Table 1). The resulting DNA template after PCR was reprecipitated and treated with the DpnI restriction enzyme. In the next step, the desired DNA template was purified on an agarose gel, phosphorylated with T4 Polynucleotide Kinase (T4 PNK), and ligated.

Treacle Δ1121-1488-NLS mutant was constructed based on the pTreacle-GFP full or pTreacle-2S full plasmid using iProof High-Fidelity DNA polymerase with primer set #7 (Supplementary Table 1). The resulting DNA template after PCR was reprecipitated and treated with the DpnI restriction enzyme. In the next step, the desired DNA template was purified on an agarose gel, phosphorylated with T4 Polynucleotide Kinase (T4 PNK), and ligated.

For the generation of Treacle ΔNoLS (Δ1350-1488) mutant, fragments were amplified by PCR from the pTreacle-GFP plasmid with primer set #8 (Supplementary Table 1). The forward and reverse primers contained BglII and BamHI sites respectively. The amplified fragment was inserted into either the pTurboGFP-C (Evrogen, FP511) or pKatushka2S-C (Evrogen, FP761) vectors using BglII/BamHI restriction/ligation.

The full-length Treacle without fluorescent protein was constructed based on the pTreacle-2S full plasmid using iProof High-Fidelity DNA polymerase with primer set #9 (Supplementary Table 1). The resulting DNA template after PCR was reprecipitated and treated with the DpnI restriction enzyme. In the next step, the desired DNA template was purified on an agarose gel, phosphorylated with T4 Polynucleotide Kinase (T4 PNK), and ligated.

Fragment of the wt Treacle (amino acids 291–426) were amplified by PCR with primer set #10 (Supplementary Table 1) and subcloned into pMAL (New England Biolabs) vector encoding the TEV protease cleavage site after MBP. cDNA fragments and the expression vector were cleaved with BamHI I and Sal I digestion and ligated under suitable conditions.

The resulting all of constructs were verified by sequencing.

### Gene knockdown

RNA interference experiments were performed using Lipofectamine 3000 transfection reagent (Thermo Scientific) following the manufacturer’s instructions. The cells were transfected with 50 nM Treacle/TCOF1 siRNA (the sequences of the siRNA is provided in Supplementary Table S3) or 50 nM TOPBP1 (Santa Cruz Biotechnology, #sc-41068).

For CRISPR/Cas9-mediated knockout, two single guide RNAs (sgRNA) to first exon of the TCOF1 gene were designed using the guide RNA design tool (www.atum.bio/eCommerce/cas9/input). The sgRNA targeting sequences were separately cloned into the px330mCherry (Addgene #98750). A list of all oligonucleotides is provided in Supplementary Table S3. The plasmids were co-transfected into HeLa cells with LTX transfection reagent (Invitrogen). 5-7 days after transfection cells were fixed and immunostained with required antibodies.

### Fluorescence microscopy

For immunostaining, cells were grown on microscope slides. All samples were fixed in 1% formaldehyde in PBS for 15 min at room temperature and treated with 1% Triton X-100 for permeabilization. Cells were washed in PBS and then incubated with antibodies in PBS supplemented with 1% BSA and 0.05% Tween 20 for 1 h at room temperature or overnight at 4°C. Then the cells were washed with PBS three times (5 min each time). The primary antibodies bound to antigens were visualized using Alexa Fluor 488-conjugated secondary antibodies. The DNA was counterstained with the fluorescent dye 4,6-diamino-2-phenylindole (DAPI) for 10 min at room temperature. The samples were mounted using Dako fluorescent mounting medium (Life Technologies). The immunostained samples were analyzed using a Zeiss AxioScope A.1 fluorescence microscope (objectives: Zeiss N-Achroplan 40 ×/0.65 and EC Plan-Neofluar 100 ×/1.3 oil; camera: Zeiss AxioCam MRm; acquisition software: Zeiss AxioVision Rel. 4.8.2; Jena, Germany) or STELLARIS 5 Leica confocal microscope (objectives: HC PL APO 63x/1.40 oil CS2). The images were processed using ImageJ software (version 1.44) and analyzed using CellProfiler software (version 3.1.5). 3D reconstruction of *xyz* confocal datasets (*z*-stacks) was performed using Leica LAS-X software. Antibodies used in the study are listed in Supplementary Table 7.

### Cell sorting

Cells were transiently transfected with required plasmids using LTX transfection reagent (Invitrogen) or immunostained with required antibodies. Cell sorting was performed using an SH800 Cell Sorter (Sony) with a laser tuned to 488 nm for green fluorescence and 561 nm for red fluorescence. Gates were set with reference to negative controls. A minimum of 2×10^6^ events was collected for ChIP or RNA extraction.

### RNA extraction and RT-qPCR

After sorting, living or immunostained cells were pelleted by centrifugation at 3000 g for 5′ at 4°C. The supernatant was discarded. Total RNA was isolated from the pellet as described (Hrvatin et al, 2014). All RNA samples were further treated with DNase I (Thermo Scientific) to remove the residual DNA. RNA (1 µg) was reverse transcribed in a total volume of 20 µl for 1 h at 42°C using 0.4 µg of random hexamer primers and 200 U of reverse transcriptase (Thermo Scientific) in the presence of 20 U of ribonuclease inhibitor (Thermo Scientific). The cDNA obtained was analyzed by quantitative polymerase chain reaction (qPCR) using the CFX96 real-time PCR detection system (Bio-Rad Laboratories). The PCRs were performed in 20 µl reaction volumes that included 50 mM Tris–HCl (pH 8.6), 50 mM KCl, 1.5 mM MgCl_2_, 0.1% Tween-20, 0.5 µM of each primer, 0.2 mM of each dNTP, 0.6 µM EvaGreen (Biotium), 0.2 U of Hot Start Taq Polymerase (Sibenzyme) and 50 ng of cDNA. Primers used in the study are listed in Supplementary Table 4.

### Chromatin immunoprecipitation (ChIP) and ChIP-seq analysis

Living cells were fixed for 15 min with 1% formaldehyde at room temperature, and crosslinking was quenched by adding 125 mM glycine for 5 min. Cell sorting was performed if needed. Cells were harvested in PBS, and nuclei were prepared by incubation in FL buffer (5 mM PIPES, pH 8.0, 85 mM KCl, 0.5% NP40) supplemented with Protease Inhibitor Cocktail (Bimake) and Phosphatase Inhibitor Cocktail (Bimake) for 30 min on ice. Next, chromatin was sonicated in RIPA buffer (10 mM Tris–HCl, pH 8.0, 140 mM NaCl, 1% Triton X-100, 0.1% sodium deoxycholate, 0.1% SDS) with a VirSonic 100 to an average length of 200-500 bp. Per ChIP reaction, 10–20 µg chromatin was incubated with 2–4 µg antibodies overnight at 4°C. The next day, Protein A/G Magnetic Beads (Thermo Scientific) were added to each sample and incubated for 4 h at 4°C. Immobilized complexes were washed two times for 10 min at 4°C in low salt buffer (20 mM Tris-HCl, pH 8.0, 150 mM NaCl, 2 mM EDTA, 0.1% SDS, 1% Triton X-100) and high salt buffer (20 mM Tris–HCl, pH 8.0, 500 mM NaCl, 2 mM EDTA, 0.1% SDS, 1% Triton X-100). Samples were incubated with RNase A (Thermo Scientific) for 30 min at room temperature. The DNA was eluted from the beads and de-crosslinked by proteinase K digestion for 4 h at 55°C and subsequent incubation at 65°C for 12 h. Next, DNA was purified using phenol/chloroform extraction and analyzed by qPCR. The qPCR primers used for ChIP analysis are listed in Supplementary Table 5. The sequencing libraries were then prepared with NEBNext Ultra II kit according manufacturer’s protocol. Final libraries were PCR amplificated and adapter dimers were cleaned with 1:1 MagPure magnetic beads (Magen Biotechnology). Resulted DNA was resuspended in 30 mkl 10 mM Tris-HCl buffer ph 8.0 and were sequenced on Illumina machine. Chip-seq reads were mapped to the reference human genome hg38 assembly using Bowtie v2.2.3 with the ‘–very-sensitive’ mode. Non-uniquely mapped reads, possible PCR and optical duplicates were filtered out using SAMtools v1.5. The bigWig files with the ratio of RPKM normalized ChIP-seq signal to the input were generated using deepTools v3.4.2 bamCompare function. Cool files was generated, merged and normalized using cooler version 0.8.11 (https://github.com/open2c/cooler).

### Electron microscopy

Twenty-four hours after transfection, cells were fixed with 2.5% neutralized glutaraldehyde in the requisite buffer for 2 h at room temperature, post-fixed with 1% aqueous OsO4, and embedded in Epon. Sections of 100 nm thickness were cut and counterstained with uranyl acetate and lead citrate. Sections were examined and photographed with a JEM 1400 transmission electron microscope (JEOL, Japan) equipped with a QUEMESA bottom-mounted CCD-camera (Olympus SIS, Japan) and operated at 100 kV.

### Protein purification

E. coli BL21 (DE3) cells were transformed with vectors encoding cDNAs of TCOF LCR fragments fused with TEV-cleavable maltose-binding protein (MBP). BL21 (DE3) were cultured in Terrific Broth medium at 37°C to OD=0.6-1.0. Protein expression was induced by adding 0.8 mM IPTG, and the cultures were further incubated at 18 °C overnight. The cells were collected by centrifugation (5,000g, 20 min, 4 °C). To purify MBP-tagged protein under a native condition, the cell pellet was dissolved in start buffer (20 mM Tris–HCl, 150 mM NaCl, 10 mM MgCl2, 0.1% NP-40, 10% glycerol, 1 mM DTT, 1 mM phenylmethylsulfonyl fluoride, 1x protease inhibitor cocktail (B14001 selleckchem), pH 7.4). Cells were disrupted by sonication in start buffer on ice. The lysate was centrifuged (20,000g, 4 °C, 60 min) and the supernatant was collected and applied to a column containing amylose resin (NEB, E8021S) at 4 °C. The column was washed with wash buffer (20 mM Tris–HCl, 500 mM NaCl, 10 mM MgCl2, 0.1% NP-40, 10% glycerol, 1 mM DTT, pH 7.4).

For cleavage of MBP TEV protease was added at molar ratio approximately 1:50 directly to the eluted protein in buffer (20 mM Tris–HCl, 200 mM NaCl, 5 mM sodium citrate, 1 mM DTT, pH 7.4) overnight at 4 °C while gently mixing the resin on a rotator. Subsequently, the tagged proteins were eluted and then subjected to ion-exchange chromatography with 0-1000 mM NaCl gradient.

Eluted proteins were sequentially dialysed at 4 °C against assay buffer (20 mM Tris–HCl pH 7.4) at 4 °C overnight and concentrated using Amicon centrifugal filters (Millipore) and stored at −80 °C in small aliquots.

### *In vitro* Treacle condensation analysis

Recombinant Treacle fragment (amino acids 291–426) was diluted in 20 mM Tris–HCl pH 7.4 to a final concentration of 100 µM. To assemble condensates PEG 8000 was added to the protein solution to a final concentration of 5%, followed by incubation at room temperature for 10 minutes. 10 μL of the resulting solution was pipetted onto a glass slide, covered with a coverslip, and analyzed on the Zeiss AxioScope A.1 fluorescence microscope in DIC mode. To test the effects of NaCL or 1,6-hexandiol on Treacle condensation, NaCL or 1,6-hexandiol was added to the protein solution with 5% PEG 8000 at 500 mM or 10% concentrations, respectively.

### Live-cell imaging

Cells were seeded in 35-mm glass-bottom dishes twelve hours before transfection with the plasmid. Twenty-four hours after transfection, Hoechst 33342 (Cell Signaling Technology) was added to the medium at a final concentration of 1 μg/ml, and the cells were incubated for 20 min at 37°C. Hoechst-containing medium was replaced with a complete fresh medium before the cells underwent live-cell imaging using a STELLARIS 5 Leica confocal microscope (objectives: HC PL APO 63x/1.40 oil CS2) equipped with an incubation chamber to provide a humidified atmosphere at 37°C with 5% CO_2_. For observation of liquid droplet behavior, z-stack time-lapses were taken. Maximum intensity projections are shown, but all findings were confirmed on a single plane.

### Optodroplet assay

Cells were seeded in 35-mm glass-bottom dishes and were transfected with Treacle-FusionRed-Cry2 plasmid using LTX transfection reagent (Invitrogen) 24 h before imaging. Twenty-four hours after transfection, Hoechst 33342 was added to the medium at a final concentration of 1 μg/ml, and the cells were incubated for 10 min at 37°C. Hoechst-containing medium was replaced with a complete fresh medium before the cells underwent live-cell imaging using a STELLARIS 5 Leica confocal microscope (objectives: HC PL APO 63x/1.40 oil CS2) equipped with an incubation chamber to provide a humidified atmosphere at 37°C with 5% CO2. Droplet formation was induced with light pulses at 488 nm (blue light, 1% laser power) for 10 sec, and z-stack images were captured every 60 sec in the absence of blue light.

### Fluorescence recovery after photobleaching (FRAP) analysis

Fluorescence recovery after photobleaching (FRAP) experiments were performed on a STELLARIS 5 Leica confocal microscope equipped with an HC PL APO 63x/1.40 oil CS2 objective and an incubation chamber that maintained a humidified atmosphere at 37°C with 5% CO₂. Live-cell experiments were performed using transfected cells cultured in phenol red-free DMEM medium supplemented with 10% fetal bovine serum. Imaging was typically performed at a resolution of 512 × 512 pixels, with a scan speed of 200 ms per frame.

Typically, images were acquired at 512 × 512 pixels at a scan speed corresponding to 200 ms per image. Before photobleaching, 3 to 5 images were recorded. Bleaching parameters, i.e., laser intensity and scanning time, were chosen to reach approximate 50% of bleaching in the shortest possible time. The bleaching area was selected according to the type of experiment (full-, partial- or half-FRAP). In the case of half-FRAP, which relies on the analysis of the bleached and the non-bleached half, it is important to optimize the bleaching step (intensity and area of the bleaching spot) to minimize bleaching in the other (non-bleached) half. Half-FRAP experiments were conducted until the signals in both halves had converged to each other or until the signal in the non-bleached half had reached its initial pre-bleach value. FRAP data analysis was performed as described in (Muzzopappa et al., 2022).

### Proximity ligation assay

After drug treatments or at 24 h post-transfection, cells were fixed for 15 min with 1% formaldehyde at room temperature and permeabilized with 1% Triton X-100. Next, cells were subjected to the proximity ligation assay (PLA) using the Duolink Green (Sigma-Aldrich) according to the manufacturer’s instructions. Briefly, cells were blocked, incubated with appropriate primary antibodies overnight at 4°C, and then incubated with PLA probes, which are secondary antibodies (anti-mouse-IgG and anti-rabbit-IgG) conjugated to unique oligonucleotides. Samples were then treated with a ligation solution followed by an amplification solution containing polymerase and fluorescently labeled oligonucleotides, allowing rolling-circle amplification and detection of discrete fluorescent dots. Some samples after the PLA protocol were additionally counterstained with Alexa-Fluor-488- or Alexa-Fluor-594-conjugated (Molecular Probes) secondary antibody. The DNA was counterstained with DAPI for 10 min at room temperature. The samples were mounted using Dako fluorescent mounting medium (Life Technologies) and analyzed using STELLARIS 5 Leica confocal microscope (objectives: HC PL APO 63x/1.40 oil CS2). The images were analyzed using ImageJ software (version 1.44).

### Transcription labeling

For EU incorporation, the cells were incubated with 100 μM EU (Sigma-Aldrich) for 2 hr at 37°C. Then, the cells were washed three times with PBS and fixed in ice-cold methanol for 10 min. The samples were then processed using a Click-iT EU Imaging Kit (Life Technologies) according to the manufacturer’s recommendations.

### Single molecule RNA Fluorescentin in situ Hybridization (smFISH)

All single-molecule RNA FISH probes were designed as described (Yao et al., 2019) and labeled with FITC on the 3’ends (Supplementary Table 6). Cells were fixed with 4% PFA for 15 min, followed by permeabilization with 1% Triton X-100 for 10 min. Cells were incubated in 10% formamide/2x SSC for 10 min at room temperature and then were hybridizated with 5 nM each of RNA probes in 50% formamide/2x SSC at 37◦C for 16 hours. After hybridization, the cells were washed 10% formamide/2x SSC for 30 min at 37◦C.

## Supporting information

MovieS1 (Intranucleolar Treacle-2S fusion events (low expression, AMD-treated cells)

MovieS2 (Intranucleolar Treacle-2S fusion events (low expression, VP16-treated cells)

MovieS3 (Extranucleolar Treacle-2S fusion events (high expression)

MovieS4 (Treacle-2S FCs half-FRAP)

MovieS5 Treacle-2S AMD half-FRAP

MovieS6 (Treacle-2S Intranucleolar condensates type 1 half-FRAP)

MovieS7 (Treacle-2S Intanucleolar condensate type 2 half-FRAP)

## Acknowledgments

This work was supported by RSF grant 21-74-10018. Confocal microscopy studies were supported by grant 075-15-2019-1661 from the Ministry of Science and Higher Education of the Russian Federation.

## Author contributions

O.L.K., A.K.V., and S.V.R. conceived the study. A.K.V. and O.L.K. designed the experiments. A.K.V., A.P.K., A.V.L., and D.A.D. performed most of the experiments. E.P.K. and I.I.K. performed transmission electron microscopy imaging. O.L.K. and A.K.V. wrote the manuscript with assistance from the other authors.

## Competing interests

The authors declare no competing interests.

## Data and materials availability

All data needed to evaluate the conclusions in the paper are present in the paper and/or the Supplementary Materials.

## Notes

### Competing Interest Statement

The authors have declared no competing interest.

### Summary of Updates

Below, we provide a concise summary of the key modifications and updates: FRAP experiments: We conducted additional FRAP experiments, including half-FRAP and full-FRAP analyses, which expanded the interpretation of our results on the dynamic behavior of Treacle in condensates. Treacle condensation experiments: We conducted the condensation experiments using a self-produced and purified recombinant fragment of Treacle. New chapter on Treacle partner interactions: We added a new section titled "Treacle condensation is essential for its proper interaction with nucleolar subcompartments", which examines the interactions of Treacle and its mutant forms with partner proteins. This section expands our interpretation of the role of Treacle's interactions in its condensation and the formation of fibrillar centers (FCs). Additional cell lines: We included additional experiments with normal human skin fibroblasts (to validate CRISPR/Cas9-mediated knockout of Treacle) and in MCF7 cells (to validate the etoposide-mediated rDNA damage response). These additions enhance the generalizability of our findings. Terminology adjustments: We revised the terminology throughout the manuscript, replacing the terms "liquid-phase" and "liquid" with "liquid-like" where appropriate. Additionally, we replaced "LLPS" with terms such as "condensation" or "coacervation" to reflect the nuanced nature of our findings. Expanded supporting materials: We enhanced the supporting materials by adding additional videos and wide-field images of cell fields for most observed patterns. Key changes to figures: Figure 1: Panel C has been replaced, as supplementing the experiment with necessary controls required redoing the entire experiment. Figure 2 now includes condensation experiments with the recombinant fragment of Treacle. Partial FRAP experiments have been replaced with half-FRAP analysis. Figure 3: The original Figure 3 has been divided into two separate figures (Figures 3 and 4 in the revised MS). FRAP analysis diagrams have been replaced and recalculated with new normalization. Figure 4: Supplemented with the analysis of Treacle mutant lacking the SE-rich LCR central domain. Figure 5: Newly added figure to display results on partner interactions of Treacle and its mutants. Figures 6 and 7: Correspond to Figures 4 and 5 of the original manuscript, with updated labeling and presentation.

