## Supplementary Figures for "Treacle’s ability to form liquid-like phase condensates is essential for nucleolar fibrillar center assembly, efficient rRNA transcription and processing, and rRNA gene repair"

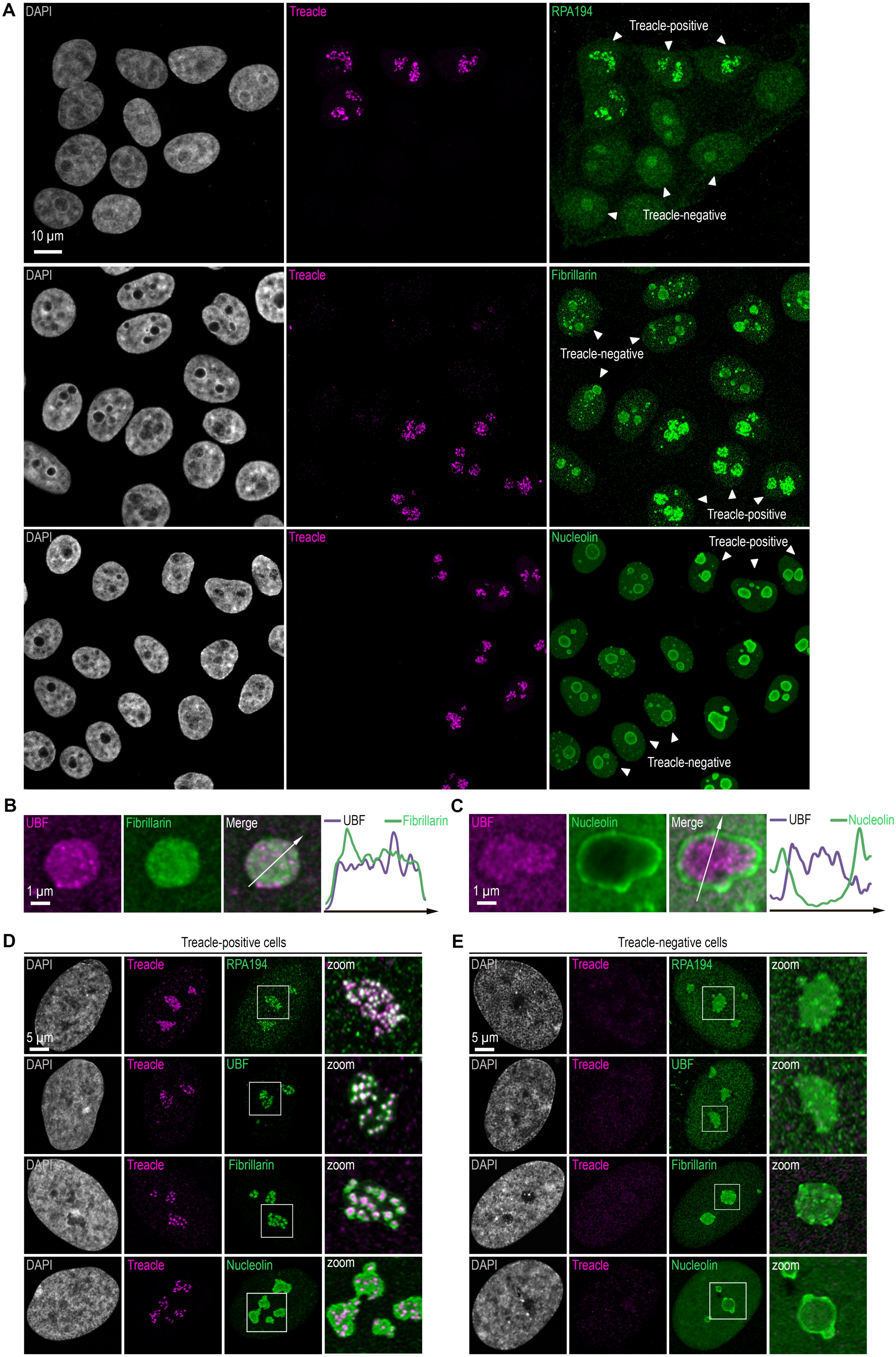

A
DAPI
Treacle
RPA194
Treacle-positive
Treacle-negative
10 μm
DAPI
Treacle
Fibrillarin
Treacle-negative
Treacle-positive
DAPI
Treacle
Nucleolin
Treacle-positive
Treacle-negative
B
UBF
Fibrillarin
Merge
UBF
Fibrillarin
1 μm
C
UBF
Nucleolin
Merge
UBF
Nucleolin
1 μm
D
Treacle-positive cells
DAPI
Treacle
RPA194
zoom
5 μm
DAPI
Treacle
UBF
zoom
DAPI
Treacle
Fibrillarin
zoom
DAPI
Treacle
Nucleolin
zoom
E
Treacle-negative cells
DAPI
Treacle
RPA194
zoom
5 μm
DAPI
Treacle
UBF
zoom
DAPI
Treacle
Fibrillarin
zoom
DAPI
Treacle
Nucleolin
zoom

**Fig. S1**

**(A)** HeLa cells were transfected with a construct coding CRISPR/Cas9 and sgRNA to the TCOF1 gene. After 7-10 days after transfection, the cells were fixed and co-immunostained with Treacle and either RPA194, Fibrillarin, or Nucleolin antibodies. The DNA was stained with DAPI (gray). Cells were analyzed by laser scanning confocal microscopy. Micrographs depicting representative cell fields are presented, with Treacle-positive and Treacle-negative cells denoted by white triangles. **(B)** HeLa cells were processed as described in (A) and co-immunostained with Fibrillarin and UBF antibodies. Cells were analyzed by laser scanning confocal microscopy. Representative images of Treacle-negative nucleolus (magnified images) are shown. **(C)** HeLa cells were processed as described in (A) and co-immunostained with Nucleolin and UBF antibodies. **(D)** Intact human skin fibroblasts (Treacle-positive) were fixed and co-immunostained with Treacle and with either RPA194, UBF, Fibrillarin or Nucleolin antibodies. DNA was stained with DAPI (gray). Cells were analyzed by laser scanning confocal microscopy. Representative images of cells and nucleoli (magnified images) are shown. **(E)** Human skin fibroblasts cells were transfected with a construct coding CRISPR/Cas9 and sgRNA to the TCOF1 gene. After 7-10 days after transfection, the cells were fixed and co-immunostained with Treacle and either RPA194, UBF, Fibrillarin or Nucleolin antibodies. DNA was stained with DAPI (gray). Cells were analyzed by laser scanning confocal microscopy. Representative images of Treacle-negative cells and nucleoli (magnified images) are shown.

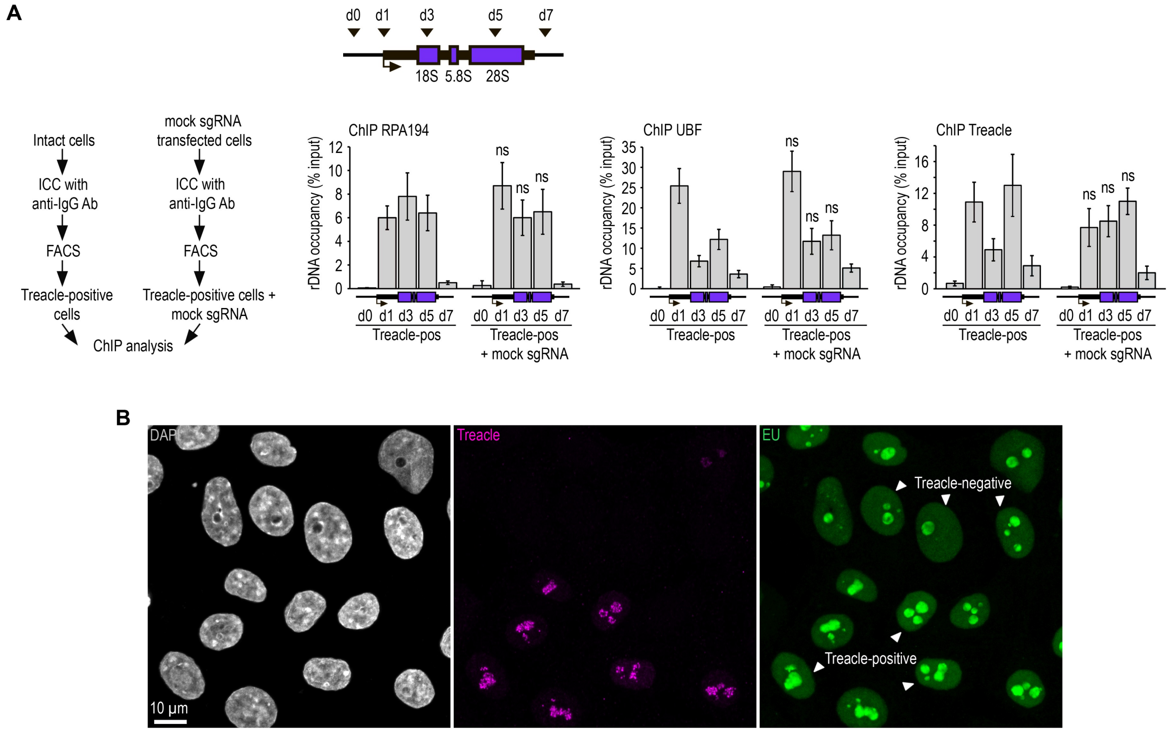


**Fig. S2**

**(A)** HeLa cells were transfected with a construct encoding CRISPR/Cas9 and a mock sgRNA. At 7–10 days post-transfection, cells were fixed, immunostained with IgG antibodies, and processed through FACS-related procedure in light scattering analysis mode (Treacle-positive cells + mock sgRNA). Similarly, intact HeLa cells were fixed, immunostained with IgG antibodies, and processed through FACS-related procedure in light scattering analysis mode (Treacle-positive cells). The resulting cell populations were utilized for ChIP analysis. ChIP was followed by qPCR using primers (d0, d1, d3, d5, d7) targeting specific regions of the rRNA gene, as indicated in the schematic. Data are presented relative to the input and expressed as means ± SD from at least three independent replicates. n.s., not significant by unpaired t test. **(B)** HeLa cells were transfected with a construct coding CRISPR/Cas9 and sgRNA to the TCOF1 gene. After 7-10 days after transfection the cells were pulsed with EU (100 μM for 2 hr), fixed and immunostained with Treacle antibodies. EU (green) was revealed by click chemistry. The DNA was stained with DAPI (gray). Cells were analyzed by laser scanning confocal microscopy. Micrographs depicting representative cell fields are presented, with Treacle-positive and Treacle-negative cells denoted by white triangles.

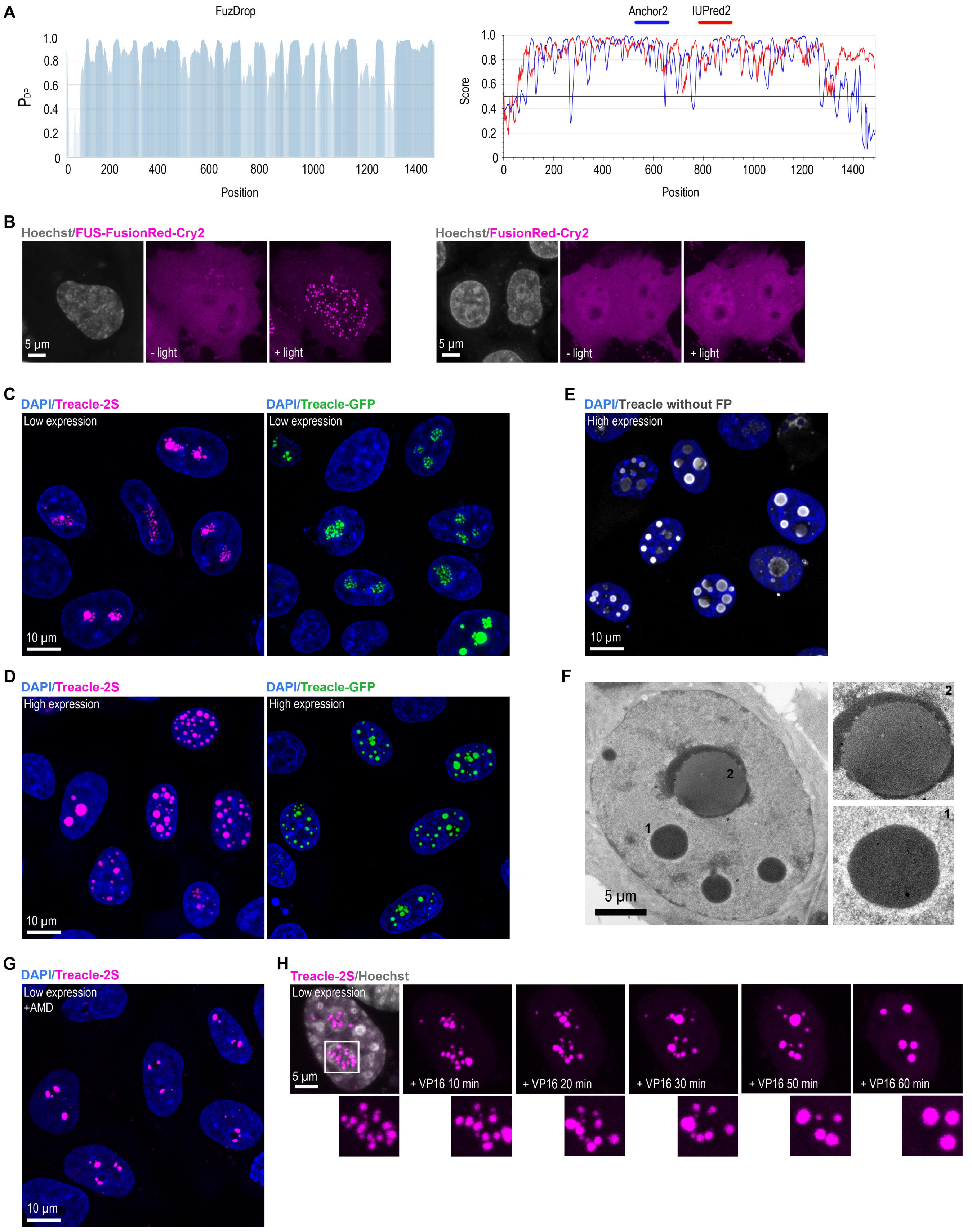
A
FuzDrop
P_DP
1.0
0.8
0.6
0.4
0.2
0
0 200 400 600 800 1000 1200 1400
Position
Anchor2
IUPred2
Score
1.0
0.8
0.6
0.4
0.2
0
0 200 400 600 800 1000 1200 1400
Position
B
Hoechst/FUS-FusionRed-Cry2
5 µm
- light
+ light
Hoechst/FusionRed-Cry2
5 µm
- light
+ light
C
DAPI/Treacle-2S
Low expression
10 µm
DAPI/Treacle-GFP
Low expression
E
DAPI/Treacle without FP
High expression
10 µm
D
DAPI/Treacle-2S
High expression
10 µm
DAPI/Treacle-GFP
High expression
F
2
1
5 µm
2
1
G
DAPI/Treacle-2S
Low expression
+AMD
10 µm
H
Treacle-2S/Hoechst
Low expression
5 µm
+ VP16 10 min
+ VP16 20 min
+ VP16 30 min
+ VP16 50 min
+ VP16 60 min

**Fig. S3**

**(A)** Full-length Treacle disorder characteristics predicted by FuzDrop, Anchor2 and IUprep2. **(B)** HeLa cells were transfected with FusionRed-Cry2 or FUS-FusionRed-Cry2 constructs and, 24 h after transfection, were illuminated with blue light for 10 sec. DNA was stained with Hoechst 33342 (gray). **(C)** HeLa cells were transfected with Treacle-Katushka2S (Treacle-2S) or Treacle-GFP at a quantity of 50 ng plasmids per $2x10^5$ cells. For low levels of expression analysis, the cells were fixed 16-24 after transfection. DNA was stained with DAPI (blue). Cells were analyzed by laser scanning confocal microscopy. Micrographs depicting representative cell fields are presented. **(D)** HeLa cells were transfected with Treacle-2S or Treacle-GFP at a quantity of 50 ng plasmids per $2x10^5$ cells. For high levels of expression analysis, the cells were fixed 48 after transfection. DNA was stained with DAPI (blue). Cells were analyzed by laser scanning confocal microscopy. Micrographs depicting representative cell fields are presented. **(E)** HeLa cells were transfected with Treacle-2S in the absence of a fused fluorescent protein (Treacle without FP). For high levels of expression analysis, the cells were fixed 48 after transfection and immunostained with Treacle antibodies. The DNA was stained with DAPI (blue). Micrographs depicting representative cell fields are presented. **(F)** HeLa cells were transfected with Treacle-2S and, 24 h after transfection were fixed and subjected to transmission electron microscopy imaging. **(G)** 16-24 h after transfection with Treacle-2S, HeLa cells were treated with 0.05μg/ml actinomycin D (AMD) to induce rDNA transcriptional repression and subsequent nucleolar cap formation. The cells were fixed and the DNA was stained with DAPI (blue). Micrographs depicting representative cell fields are presented. **(H)** HeLa cells were transfected with Treacle-2S. 24 h after transfection, the cells were treated with 90 μM VP16 to induce DNA breaks and live time-lapse imaged for 60 minutes. Representative examples of Treacle-2S fusions (magnified regions) are shown. DNA was stained with Hoechst 33342 (gray).

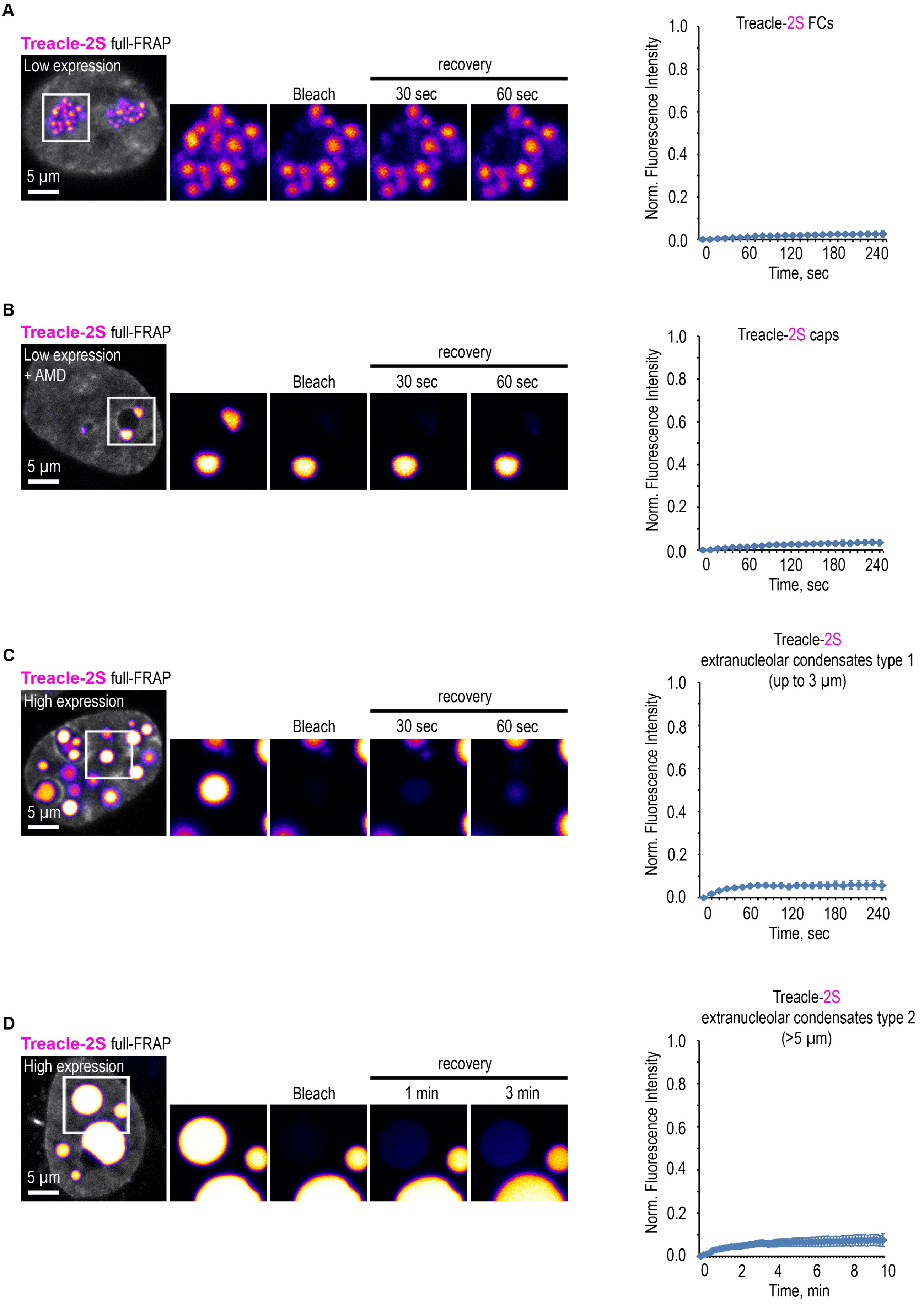
A
Treacle-2S full-FRAP
Low expression
5 µm
Bleach
recovery
30 sec
60 sec
Treacle-2S FCs
Norm. Fluorescence Intensity
Time, sec
B
Treacle-2S full-FRAP
Low expression
+ AMD
5 µm
Bleach
recovery
30 sec
60 sec
Treacle-2S caps
Norm. Fluorescence Intensity
Time, sec
C
Treacle-2S full-FRAP
High expression
5 µm
Bleach
recovery
30 sec
60 sec
Treacle-2S
extranucleolar condensates type 1
(up to 3 µm)
Norm. Fluorescence Intensity
Time, sec
D
Treacle-2S full-FRAP
High expression
5 µm
Bleach
recovery
1 min
3 min
Treacle-2S
extranucleolar condensates type 2
(>5 µm)
Norm. Fluorescence Intensity
Time, min

**Fig. S4**

**(A)** HeLa cells were transiently transfected with Treacle-2S. 16-24 h after transfection full-FRAP analysis of Treacle-labelled fibrillar center (FCs) was performed. One of the FC was bleached, and fluorescence recovery was monitored. Representative time-lapse images of the photobleached FC are shown (magnified images). DNA was stained with Hoechst 33342 (gray). Graph illustrate the quantification of the FCs full-FRAP analysis. Each trace represents an average of measurements for at least twenty FCs; error bars represent SD. **(B)** 16-24 h after transfection with Treacle-2S, HeLa cells were treated with 0.05µg/ml AMD to induce the formation of nucleolar caps. Full-FRAP analysis of Treacle-labelled nucleolar caps was performed. Representative time-lapse images of photobleached one of the nucleolar cap are shown (magnified images). DNA was stained with Hoechst 33342 (gray). Graph illustrate the quantification of the nucleolar caps full-FRAP analysis. Each trace represents an average of measurements for at least twenty caps; error bars represent SD. **(C)** HeLa cells were transiently transfected with Treacle-2S. 48 h after transfection, full-FRAP analysis of Treacle-2S extranucleolar condensates with diameters of up to 3 µm (type 1) was performed. One of the condensate of the first type was bleached, and fluorescence recovery was monitored. Representative time-lapse images of photobleached Treacle-2S extranucleolar condensates of the first type are shown (magnified images). DNA was stained with Hoechst 33342 (gray). Graph illustrate the quantification of the extranucleolar condensates full-FRAP analysis. Each trace represents an average of measurements for at least twenty extranucleolar condensates of the first type; error bars represent SD. **(D)** Full-FRAP analysis of Treacle-2S extranucleolar condensates exceeding 5 µm in diameter (type 2) was performed as described in (C).

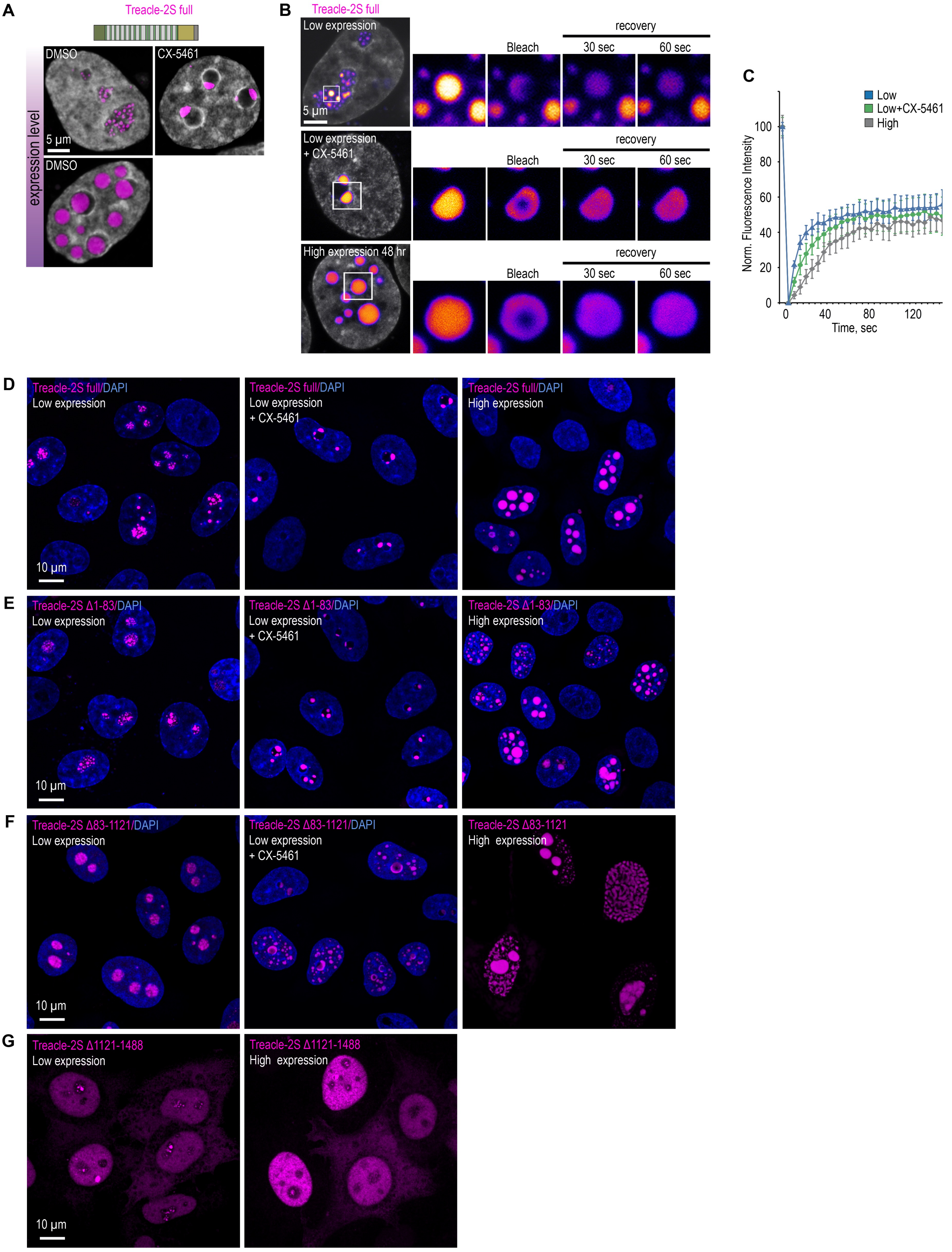
A
Treacle-2S full
DMSO
CX-5461
5 μm
DMSO
expression level
B
Treacle-2S full
Low expression
5 μm
Bleach
recovery
30 sec
60 sec
Low expression
+ CX-5461
High expression 48 hr
C
Low
Low+CX-5461
High
Norm. Fluorescence Intensity
Time, sec
D
Treacle-2S full/DAPI
Low expression
Low expression
+ CX-5461
High expression
10 μm
E
Treacle-2S Δ1-83/DAPI
F
Treacle-2S Δ83-1121/DAPI
Treacle-2S Δ83-1121
G
Treacle-2S Δ1121-1488
Low expression
High expression

**Fig. S5**

**(A)** HeLa cells were transfected with full-length Treacle-2S. For low or high levels expression analysis cells were cultivated 16-24 or 48 h after transfection respectively. The expression level is indicated by the colored zone to the left of the cell images. HeLa cells with low expression level were additionally treated with CX-5461 to induce rDNA transcriptional repression and subsequent nucleolar cap formation. Cells were fixed and analyzed by laser scanning confocal microscopy. DNA was stained with DAPI (blue). Representative images of full-length Treacle-2S condensate are shown. **(B)** HeLa cells were transfected with full-length Treacle-2S and processed as described in (A). Partial FRAP analysis of full-length Treacle-2S condensates was performed. A part of each condensate type was photobleached, and the subsequent fluorescence recovery was monitored. Representative time-lapse images of the photobleached condensates are shown (magnified images). DNA was stained with Hoechst 33342 (gray). **(C)** HeLa cells were transfected with full-length Treacle-2S and processed as described in (A). Partial FRAP analysis of full-length Treacle-2S condensates was performed as described in (B). Graphs illustrate the quantification of the full-length Treacle-2S condensates partial FRAP analysis. Each trace represents an average of measurements for at least twenty full-length Treacle-2S condensates of each type; error bars represent SD. **(D)** HeLa cells were transfected with full-length Treacle-2S and processed as described in (A). DNA was stained with DAPI (blue). Cells were analyzed by laser scanning confocal microscopy. Micrographs depicting representative cell fields are presented. **(E)** HeLa cells were transfected with Treacle-2S Δ1-83 deletion mutant and processed as described in (A). DNA was stained with DAPI (blue). Cells were analyzed by laser scanning confocal microscopy. Micrographs depicting representative cell fields are presented. **(F)** HeLa cells were transfected with Treacle-2S Δ83-1121 deletion mutant and processed as described in (A). DNA was stained with DAPI (blue). Cells were analyzed by laser scanning confocal microscopy. Micrographs depicting representative cell fields are presented. **(G)** HeLa cells were transfected with Treacle-2S Δ1121-1488 deletion mutant fused with nuclear localization signal (NLS) from simian virus 40 (SV40). For low or high levels expression analysis cells were cultivated 16-24 or 48 h after transfection respectively. Cells were fixed and analyzed by laser scanning confocal microscopy. Representative images of cells are shown.

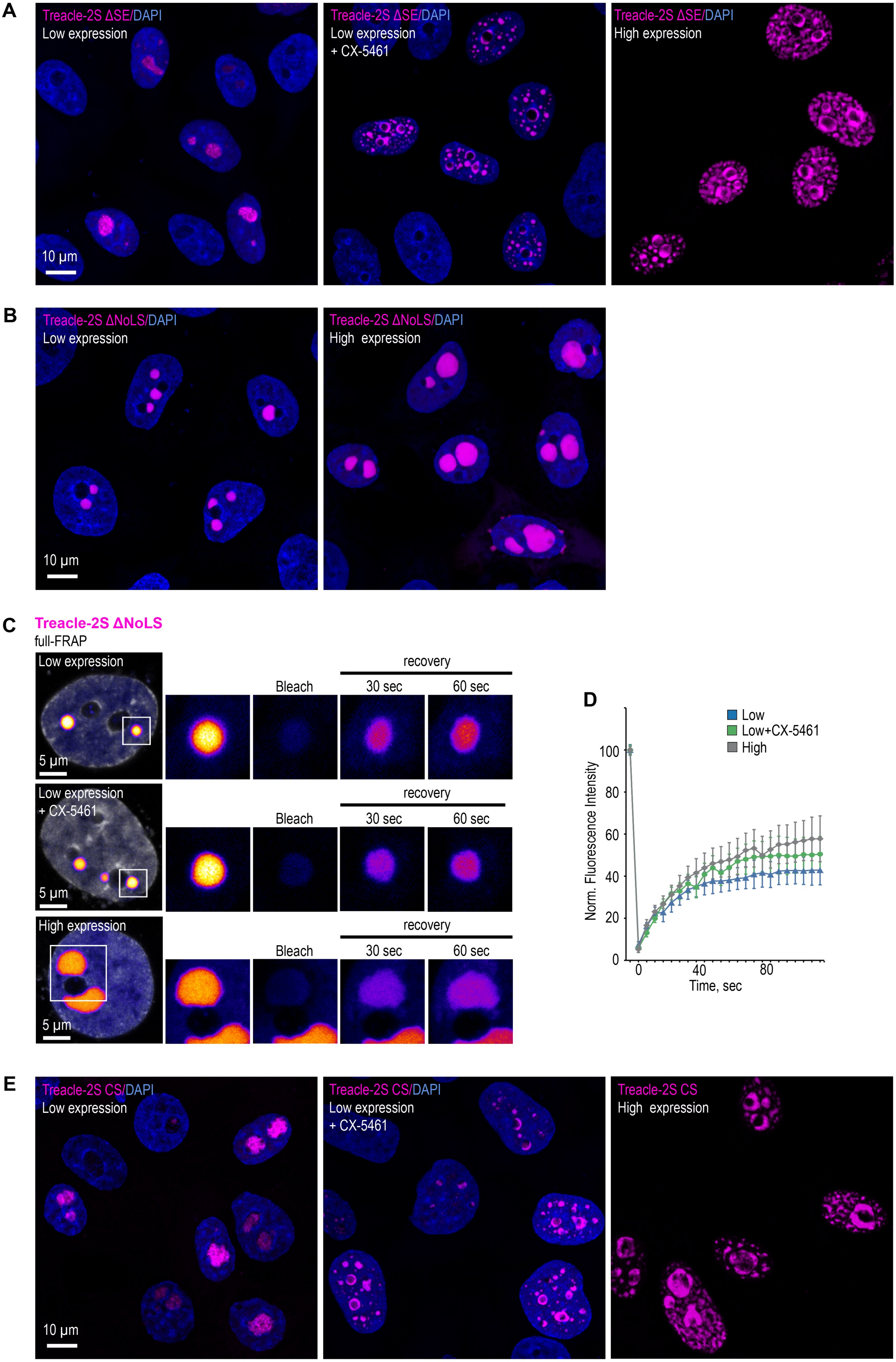
A
Treacle-2S ΔSE/DAPI
Low expression
10 µm
Treacle-2S ΔSE/DAPI
Low expression
+ CX-5461
Treacle-2S ΔSE/DAPI
High expression
B
Treacle-2S ΔNoLS/DAPI
Low expression
10 µm
Treacle-2S ΔNoLS/DAPI
High expression
C
Treacle-2S ΔNoLS
full-FRAP
Low expression
5 µm
Bleach
recovery
30 sec
60 sec
Low expression
+ CX-5461
5 µm
High expression
5 µm
D
Low
Low+CX-5461
High
Norm. Fluorescence Intensity
0
20
40
60
80
100
0
40
80
Time, sec
E
Treacle-2S CS/DAPI
Low expression
10 µm
Treacle-2S CS/DAPI
Low expression
+ CX-5461
Treacle-2S CS
High expression

**Fig. S6**

**(A)** HeLa cells were transfected with Treacle-2S ΔSE mutant. For low or high levels expression analysis cells were cultivated 16-24 or 48 h after transfection respectively. The expression level is indicated by the colored zone to the left of the cell images. HeLa cells with low expression level were additionally treated with CX-5461 to induce rDNA transcriptional repression. Cells were fixed and analyzed by laser scanning confocal microscopy. DNA was stained with DAPI (blue). Micrographs depicting representative cell fields are presented. **(B)** HeLa cells were transfected with Treacle-2S Δ1350-1488 deletion mutant (Treacle-2S ΔNoLS). For low or high levels expression analysis cells were cultivated 16-24 or 48 h after transfection respectively. Cells were fixed and analyzed by laser scanning confocal microscopy. DNA was stained with DAPI (blue). Micrographs depicting representative cell fields are presented. **(C)** HeLa cells were transiently transfected with Treacle-2S ΔNoLS. For low or high levels expression analysis cells were cultivated 16-24 or 48 h after transfection respectively. HeLa cells with low expression level were additionally treated with CX-5461. Full-FRAP analysis of Treacle-2S ΔNoLS condensates was performed. Each type of condensate was fully photobleached, and the subsequent fluorescence recovery was monitored. Representative time-lapse images of the photobleached condensates are shown (magnified images). DNA was stained with Hoechst 33342 (gray). **(D)** HeLa cells were transfected with Treacle-2S ΔNoLS mutant and processed as described in (C). Graphs illustrate the quantification of the Treacle-2S ΔNoLS condensates partial FRAP analysis. Each trace represents an average of measurements for at least twenty Treacle-2S ΔNoLS condensates of each type; error bars represent SD. **(E)** HeLa cells were transfected with charge-scrambled Treacle-2S mutant (Treacle-2S CS) and processed as described in (A).

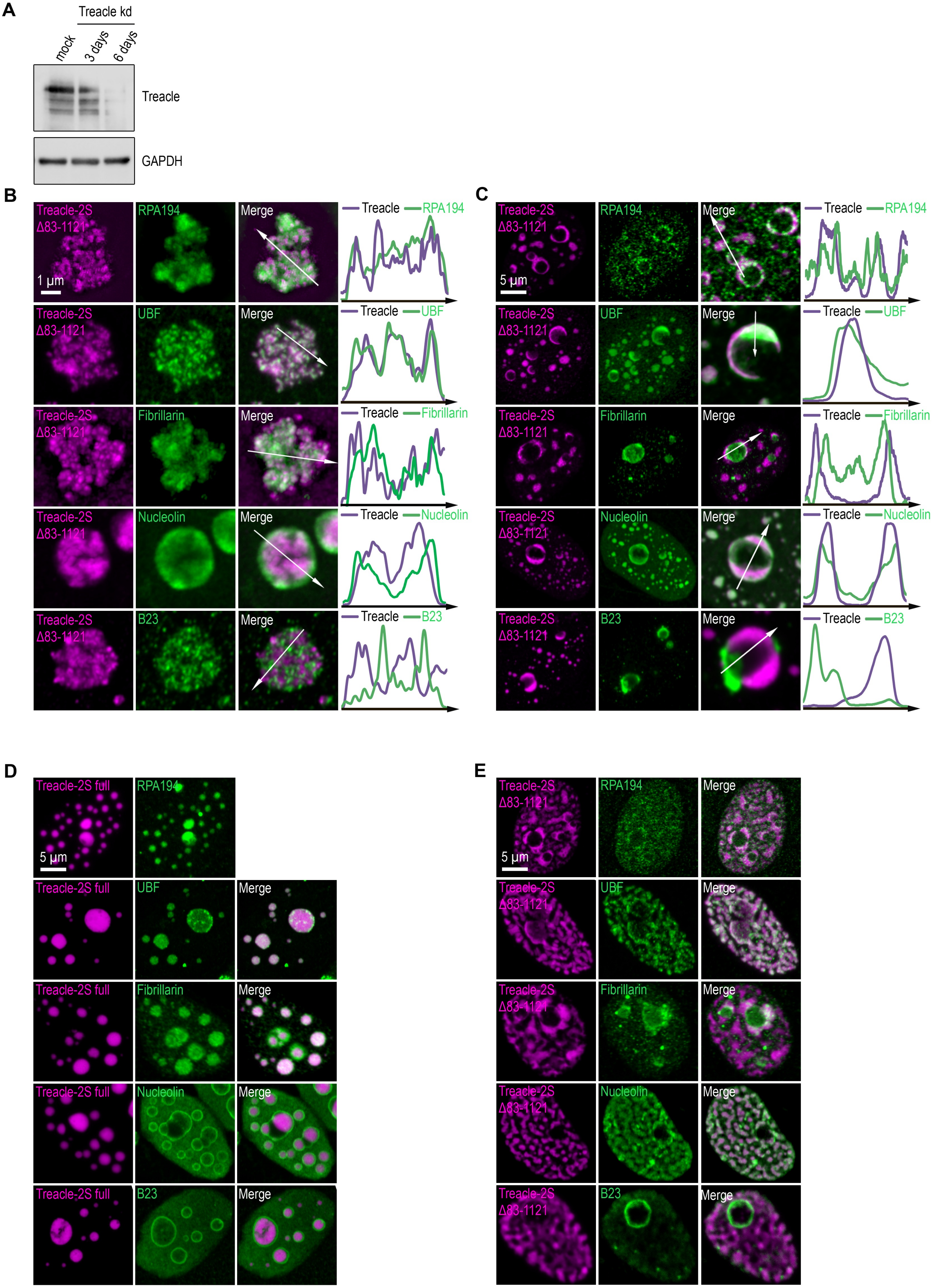
A
Treacle kd
mock
3 days
6 days
Treacle
GAPDH
B
Treacle-2S Δ83-1121
RPA194
Merge
1 μm
Treacle
RPA194
UBF
Fibrillarin
Nucleolin
B23
C
5 μm
D
Treacle-2S full
E

**Fig. S7**

**(A)** Endogenous Treacle was depleted by siRNA-mediated knockdown (Treacle kd). After 3-6 days, knockdown efficiencies were analyzed by Western blotting. mock, scrambled control siRNA. **(B)** Endogenous Treacle was depleted by siRNA-mediated knockdown (Treacle kd). Next, Treacle-depleted cells were transfected with siRNA-resistant Treacle-2S Δ83-1121 deletion mutant (Treacle-2S Δ83-1121). Cells were fixed 16-24 after transfection and immunostained with either RPA194, UBF, Fibrillarin, B23 or Nucleolin antibodies. Cells were analyzed by laser scanning confocal microscopy. Representative images of magnified nucleoli are shown. Co-localization analysis was performed on the merged images. Graphs illustrate quantification in arbitrary units of Treacle-2S variants and RPA194, UBF, Fibrillarin, B23 or Nucleolin fluorescence distribution along the lines shown in the figures. **(C)** Endogenous Treacle was depleted by siRNA-mediated knockdown (Treacle kd). Next, Treacle-depleted cells were transfected with siRNA-resistant Treacle-2S Δ83-1121 deletion mutant (Treacle-2S Δ83-1121). 16-24 h after transfection cells were treated with 0.05μg/ml actinomycin D (AMD) to induce rDNA transcriptional repression, fixed and processed as described in (B). **(D)** HeLa cells were transiently transfected with full-length Treacle-2S (Treacle-2S full). 48 h after transfection cells were fixed and immunostained with either RPA194, UBF, Fibrillarin, B23 or Nucleolin antibodies. Cells were analyzed by laser scanning confocal microscopy. Representative images of cells are shown. **(E)** HeLa cells were transiently transfected with Treacle-2S Δ83-1121 deletion mutant (Treacle-2S Δ83-1121) and processed as described in (D).

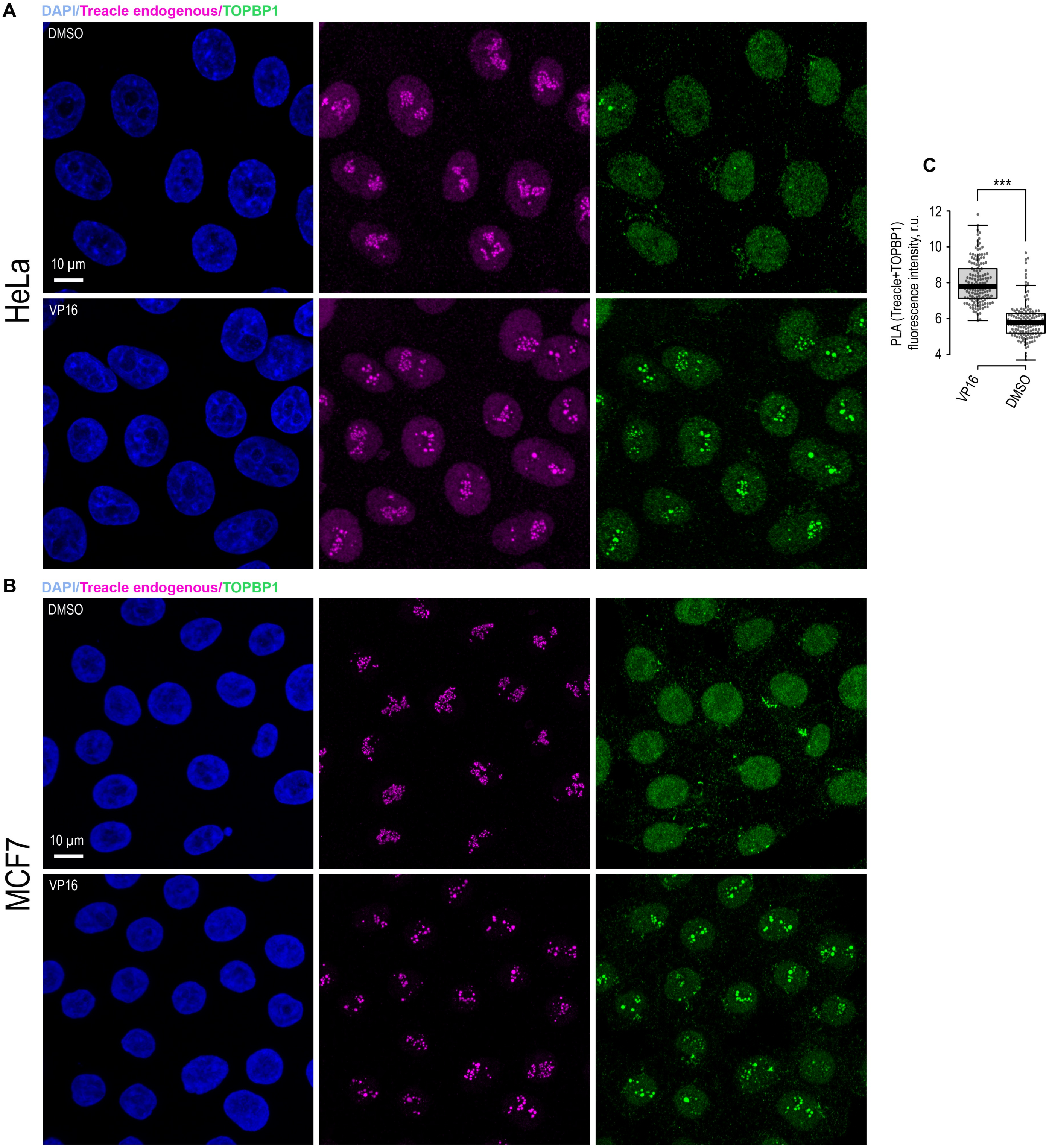


**Fig. S8**

**(A)** DMSO-treated and VP16-treated (90 µM, 30 min) HeLa cells were co-immunostained for Treacle (Treacle endogenous; magenta) and TOPBP1 (green) and analyzed by laser scanning confocal microscopy. The DNA was stained with DAPI (blue). Micrographs depicting representative cell fields are presented. **(B)** DMSO-treated and VP16-treated (90 µM, 30 min) MCF7 cells were co-immunostained for Treacle (magenta) and TOPBP1 (green) and analyzed by laser scanning confocal microscopy. The DNA was stained with DAPI (blue). Micrographs depicting representative cell fields are presented. **(C)** DMSO-treated and VP16-treated (90 µM, 30 min) HeLa cells were subjected to Proximity Ligation Assay (PLA) with antibodies against TOPBP1 and Treacle. Graphs illustrate quantification of the intensity of the PLA fluorescent signal per nucleolus (n>100).

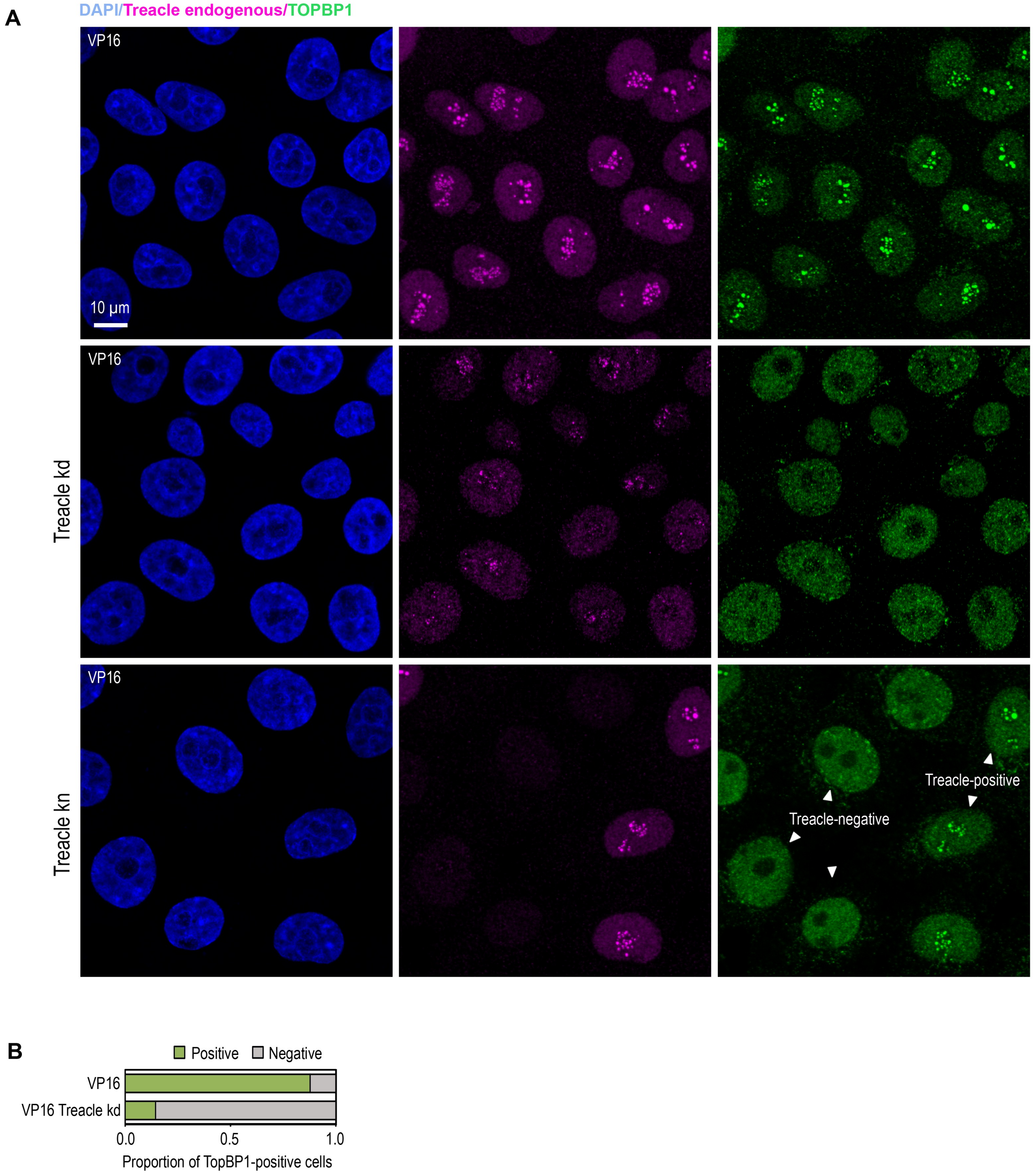


**Fig. S9**

**(A)** Intact HeLa cells, siRNA-depleted for Treacle (Treacle kd) cells or CRISPR/Cas9-depleted for Treacle (Treacle kn) cells were treated with 90 µM VP16 for 30 min. Cells were co-immunostained for TOPBP1 (green) and Treacle (magenta) antibodies and analyzed by laser scanning confocal microscopy. The DNA was stained with DAPI (blue). Micrographs depicting representative cell fields are presented. **(B)** Intact HeLa cells and Treacle kd cells were treated with 90 µM VP16 for 30 min. Cells were immunostained for TOPBP1. Percentage of cells containing TOPBP1 (TOPBP1-positive) foci within nucleoli is shown.

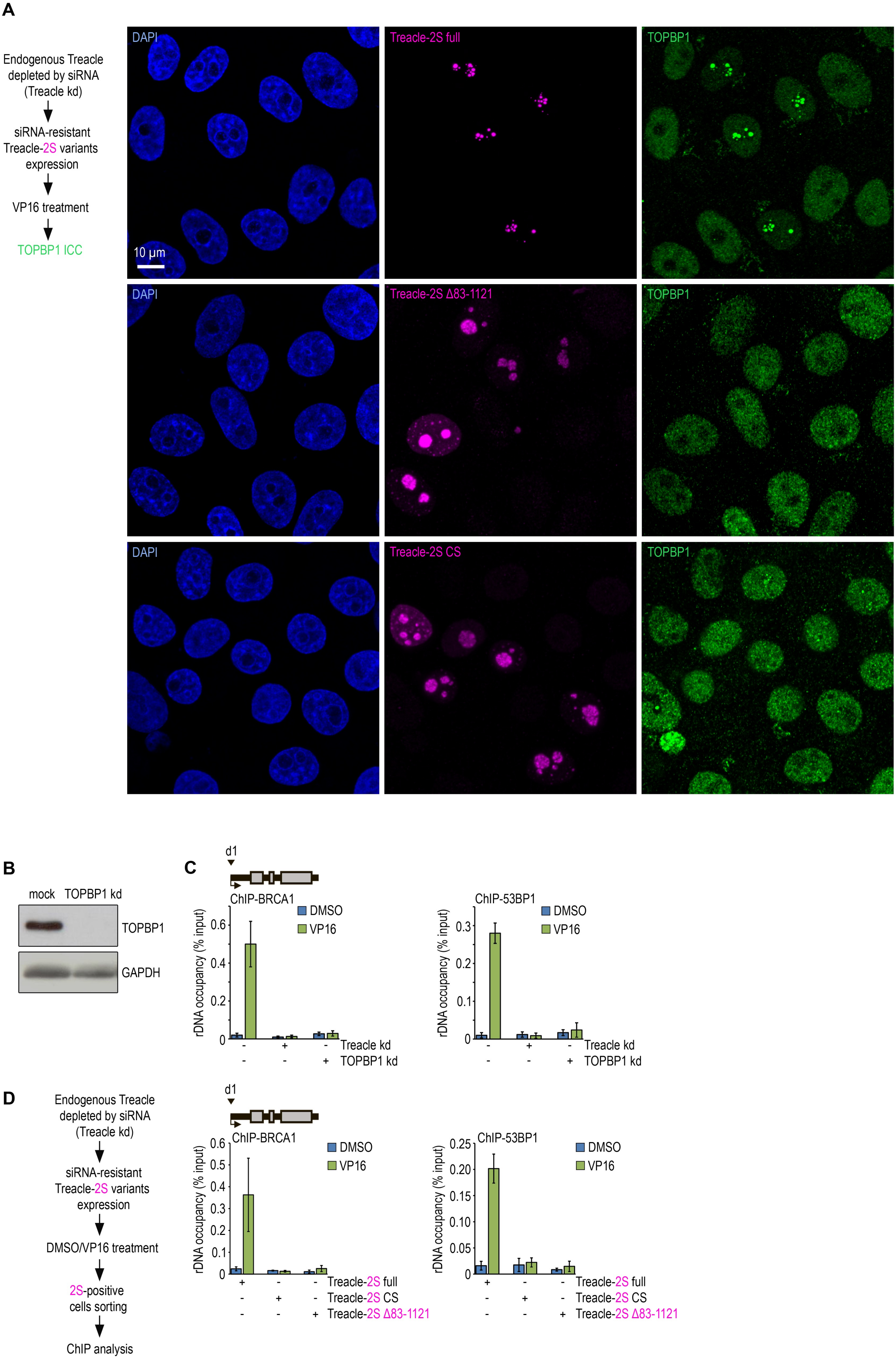
A
Endogenous Treacle depleted by siRNA (Treacle kd)
siRNA-resistant Treacle-2S variants expression
VP16 treatment
TOPBP1 ICC
DAPI
Treacle-2S full
TOPBP1
10 μm
DAPI
Treacle-2S Δ83-1121
TOPBP1
DAPI
Treacle-2S CS
TOPBP1
B
mock
TOPBP1 kd
TOPBP1
GAPDH
C
d1
ChIP-BRCA1
DMSO
VP16
rDNA occupancy (% input)
0.6
0.4
0.2
0
ChIP-53BP1
0.3
0.2
0.1
Treacle kd
TOPBP1 kd
D
Endogenous Treacle depleted by siRNA (Treacle kd)
siRNA-resistant Treacle-2S variants expression
DMSO/VP16 treatment
2S-positive cells sorting
ChIP analysis
0.5
0.3
0.1
0.25
0.20
0.15
0.10
0.05
Treacle-2S full
Treacle-2S CS
Treacle-2S Δ83-1121

**Fig. S10**

**(A)** Endogenous Treacle was depleted by siRNA-mediated knockdown (Treacle kd). Next, Treacle-depleted cells were transfected with plasmid constructs encoding either siRNA-resistant full-length Treacle-2S (Treacle-2S full), Treacle-2S Δ83-1121 deletion mutant, or charge-scrambled mutant Treacle-2S (Treacle-2S CS). 24 h after transfection, cells were treated with VP16 (90 µM for 30 min), fixed and immunostained for TOPBP1 (green) and analyzed by laser scanning confocal microscopy. The DNA was stained with DAPI (blue). Micrographs depicting representative cell fields are presented. **(B)** HeLa cells were depleted for TOPBP1 using siRNA-mediated knockdown (TOPBP1 kd). **(C)** Intact HeLa cells and cells siRNA-depleted for either Treacle (Treacle kd) or TOPBP1 (TOPBP1 kd) were treated with DMSO or 90 µM VP16 for 30 min. ChIP experiments were performed with antibodies against BRCA1 or 53BP1 antibodies. ChIP was followed by qPCR using the d1 primers to the promoter of the rRNA gene (positioned as indicated on the scheme). Data are represented relative to the input. Values are means ±SD from at least three independent replicates. **(D)** Endogenous Treacle was depleted by siRNA-mediated knockdown (Treacle kd). Treacle-depleted cells were transfected with plasmid constructs encoding siRNA-resistant either Treacle-2S full, Treacle-2S Δ83-1121 or Treacle-2S CS. 24 h after transfection, cells were treated with DMSO or VP16 (90 µM for 30 min) and fixed. Cells were subjected to cell sorting in the fluorescent analysis mode to obtain 2S-positive populations. At least $2x10^6$ sorted cells were used for ChIP with BRCA1 or 53BP1 antibodies. ChIP was followed by qPCR using the d1 primers to the promoter of the rRNA gene (positioned as indicated on the scheme). Data are represented relative to the input. Values are means ±SD from at least three independent replicates.
