## Supplementary tables 2 and 7 for "Treacle’s ability to form liquid-like phase condensates is essential for nucleolar fibrillar center assembly, efficient rRNA transcription and processing, and rRNA gene repair"

**Supplementary Table 2.**

Original Treacle full sequence

MAEARKRRELLPLIYHHLLRAGYVRAAREVKEQSGQKCFLAQPVTLLDIYTHWQQTSELGRKRKAEEDAALQAKKTRVSDPISTSESSEEEEEAEAETAKATPRLASTNSSVLGADLPSSMKEKAKAETEKAGKTGNSMPHPATGKTVANLLSGKSPRKSAEPSANTTLVSETEEEGSVPAFGAAAKPGMVSAGQADSSSEDTSSSSDETDVEGKPSVKPAQVKASSVSTKESPARKAAPAPGKVGDVTPQVKGGALPPAKRAKKPEEESESSEEGSESEEEAPAGTRSQVKASEKILQVRAASAPAKGTPGKGATPAPPGKAGAVASQTKAGKPEEDSESSSEESSDSEEETPAAKALLQAKASGKTSQVGAASAPAKESPRKGAAPAPPGKTGPAVAKAQAGKREEDSQSSSEESDSEEEAPAQAKPSGKAPQVRAASAPAKESPRKGAAPAPPRKTGPAAAQVQVGKQEEDSRSSSEESDSDREALAAMNAAQVKPLGKSPQVKPASTMGMGPLGKGAGPVPPGKVGPATPSAQVGKWEEDSESSSEESSDSSDGEVPTAVAPAQEKSLGNILQAKPTSSPAKGPPQKAGPVAVQVKAEKPMDNSESSEESSDSADSEEAPAAMTAAQAKPALKIPQTKACPKKTNTTASAKVAPVRVGTQAPRKAGTATSPAGSSPAVAGGTQRPAEDSSSSEESDSEEEKTGLAVTVGQAKSVGKGLQVKAASVPVKGSLGQGTAPVLPGKTGPTVTQVKAEKQEDSESSEEESDSEEAAASPAQVKTSVKKTQAKANPAAARAPSAKGTISAPGKVVTAAAQAKQRSPSKVKPPVRNPQNSTVLARGPASVPSVGKAVATAAQAQTGPEEDSGSSEEESDSEEEAETLAQVKPSGKTHQIRAALAPAKESPRKGAAPTPPGKTGPSAAQAGKQDDSGSSSEESDSDGEAPAAVTSAQVIKPPLIFVDPNRSPAGPAATPAQAQAASTPRKARASESTARSSSSESEDEDVIPATQCLTPGIRTNVVTMPTAHPRIAPKASMAGASSSKESSRISDGKKQEGPATQVSKKNPASLPLTQAALKVLAQKASEAQPPVARTQPSSGVDSAVGTLPATSPQSTSVQAKGTNKLRKPKLPEVQQATKAPESSDDSEDSSDSSSGSEEDGEGPQGAKSAHTLGPTPSRTETLVEETAAESSEDDVVAPSQSLLSGYMTPGLTPANSQASKATPKLDSSPSVSSTLAAKDDPDGKQEAKPQQAAGMLSPKTGGKEAASGTTPQKSRKPKKGAGNPQASTLALQSNITQCLLGQPWPLNEAQVQASVVKVLTELLEQERKKVVDTTKESSRKGWESRKRKLSGDQPAARTPRSKKKKKLGAGEGGEASVSPEKTSTTSKGKAKRDKASGDVKEKKGKGSLGSQGAKDEPEEELQKGMGTVEGGDQSNPKSKKEKKKSDKRKKDKEKKEKKKKAKKASTKDSESPSQKKKKKKKKTAEQTV

The replaced sequence is highlighted in yellow.

Insert Treacle ΔSE sequence

NSMPHPATGKTVANLLSGKSPRKSAEPSANTTLVSETEEEGSVPAFGAAAKPGMVSAGQADSSSEDTSSSSDETDVEGKPSVKPAQVKASSVSTKESPARKAAPAPGKVGDVTPQVKGGALPPAKRAKKPEEESESSEEGSESEEEAPAGTRSQVKASEKILQVRAASAPAKGTPGKGATPAPPGKAGAVASQTKAGKPEEDSESSSEESSDSEEETPAAKALLQAKASGKTSQVGAASAPAKESPRKGAAPAPPGKTGPAVAKAQAGKREEDSQSSSEESDSEEEAPAQAKPSGKAPQVRAASAPAKESPRKGAAPAPPRKTGPAAAQVQVGKQEEDSRSSSEESDSDREALAAMNAAQVKPLGKSPQVKPASTMGMGPLGKGAGPVPPGKVGPATPSAQVGKWEEDSESSSEESSDSSDGEVPTAVAPAQEKSLGNILQAKPTSSPAKGPPQKAGPVAVQVKAEKPMDNSESSEESSDSADSEEAPAAMTAAQAKPALKIPQTKACPKKTNTTASAKVAPVRVGTQAPRKAGTATSPAGSSPAVAGGTQRPAEDSSSSEESDSEEEKTGLAVTVGQAKSVGKGLQVKAASVPVKGSLGQGTAPVLPGKTGPTVTQVKAEKQEDSESSEEESDSEEAAASPAQVKTSVKKTQAKANPAAARAPSAKGTISAPGKVVTAAAQAKQRSPSKVKPPVRNPQNSTVLARGPASVPSVGKAVATAAQAQTGPEEDSGSSEEESDSEEEAETLAQVKPSGKTHQIRAALAPAKESPRKGAAPTPPGKTGPSAAQAGKQDDSGSSSEESDSDGEAPAAVTSAQVIKPPLIFVDPNRSPAGPAATPAQAQAASTPRKARASESTARSSSSESEDEDVIPATQCLTPGIRTNVVTMPTAHPRIAPKASMAGASSSKESSRISDGKKQEGPATQVSKKNPASLPLTQAALKVLAQKASEAQPPVARTQPSSGVDSAVGTLPATSPQSTSVQAKGTNKLRKPKL

The deleted SE residues

Insert Treacle CS sequence

NSMPHPATGETVANLLSGKSPEKSAEPSANTTLVSRTREEGSVPAFGAAAEPGMVSAGQADSSSRDTSSSSDRTDVEVEASEKILQVRAASAPAEGTPGKGATPAPPGEAGAVASQTKAGKPERDSESSSERSSDSRRETPAAKALLQAEASGKTSQVGAASAPAEESPRKGAAPAPPGETGPAVAKAQAGKRERDSQSSSRESDSRREAPAQAEPSGKAPQVRAASAPAEESPRKGAAPAPPRETGPAAAQVQVGKQERDSRSSSRESDSDREALAAMNAAQVEPLGKSPQVKPASTMGMGPLGKGAGPVPPGEVGPATPSAQVGKWERDSRSSSRESSDSSDGEVPTAVAPAQEESLGNILQAKPTSSPAEGPPQEAGPVAVQVKAEKPMDNSRSSRESSDSADSREAPAAMTAAQAEPALKIPQTEACPKKTNTTASAEVAPVRVGTQAPREAGTATSPAGSSPAVAGGTQRPAEDSSSSRESDSREEKTGLAVTVGQAESVGKGLQVKAASVPVEGSLGQGTAPVLPGETGPTVTQVKAEKQEDSRSSREESDSREAAASPAQVETSVKKTQAKANPAAAEAPSAKGTISAPGEVVTAAAQAKQRSPSKVEPPVRNPQNSTVLARGPASVPSVGEAVATAAQAQTGPERDSGSSREESDSRREAETLAQVEPSGKTHQIRAALAPAEESPRKGAAPTPPGETGPSAAQAGKQDDSGSSSRRSDSDGEAPAAVTSAQVIEPPLIFVDPNRSPAGPAATPAQAQAASTPREARASRSTARSSSSESRDEDVIPATQCLTPGIRTNVVTMPTAHPRIAPEASMAGASSSKESSRISDGKKQEGPATQVSEKNPASLPLTQAALKVLAQKASEAQPPVARTQPSSGVDSAVGTLPATSPQSTSVQAKGTNELRKPKL

The replaced E residues is highlighted in red, the replaced K residues is highlighted in blue.

| **Supplementary Table 7**: Antibodies used in the study | | | |
| --- | --- | --- | --- |
| **Antibody** | **Source** | **Cat. #** | **Applications** |
| anti-TopBP1, mouse | Santa Cruz Biotechnology | sc-271043 | ICC, ChIP |
| anti-B23, mouse | Sigma-Aldrich | B0556 | ICC |
| anti-pATR (Thr1989), rabbit | Cell Signaling | 58014 | ChIP |
| anti-TurboGFP, rabbit | Eurogene | AB513 | ChIP |
| anti-Katushka2S, rabbit | Eurogene | AB233 | ChIP |
| anti-53BP1, rabbit | Santa Cruz Biotechnology | sc-22760 | ICC, ChIP |
| anti-Treacle/TCOF1, mouse | Santa Cruz Biotechnology | sc-374536 | ICC |
| anti-Treacle/TCOF1, rabbit | Sigma-Aldrich | HPA038237 | ICC |
| Anti-Fibrillarin, rabbit | Abcam | ab166630 | ICC |
| Anti-RPA194, mouse | Santa Cruz Biotechnology | sc-48385 | ICC, ChIP |
| Anti-UBF1, rabbit | Thermo | PA5-82245 | ICC, ChIP |
| Anti-Nucleolin, rabbit | Cell Signaling | 14574 | ICC |
| anti-BRCA1, mouse | Santa Cruz Biotechnology | sc-6954 | ICC, ChIP |
| anti-pATM (Ser1981), mouse | Cell Signaling | 4526 | ICC, ChIP |
| anti-γH2AX (Ser139), mouse | Millipore | 05-636 | ICC**,** ChIP |
| *Abbreviations: ICC, immunocytochemistry; ChIP, chromatin immunoprecipitation;* | | | |
